# Adaptation of the yeast gene knockout collection is near-perfectly predicted by fitness and diminishing return epistasis

**DOI:** 10.1101/2022.03.25.485769

**Authors:** Karl Persson, Simon Stenberg, Markus J. Tamás, Jonas Warringer

**Affiliations:** Department of Chemistry and Molecular Biology, University of Gothenburg, PO Box 462, 40530 Gothenburg, Sweden; Department of Biology and Biological Engineering, Chalmers University of Technology, Gothenburg, Sweden

**Keywords:** Adaptation, evolvability, epistasis, diminishing return, yeast

## Abstract

Adaptive evolution of clonally dividing cells and microbes is the ultimate cause of cancer and infectious diseases. The possibility of constraining the adaptation of cell populations, by inhibiting proteins that enhance their evolvability has therefore attracted substantial interest. However, our current understanding of how individual genes influence the speed of adaptation is limited, partly because accurately tracking adaptation for many experimental cell populations in parallel is challenging. Here we use a high throughput artificial laboratory evolution (ALE) platform to track the adaptation of >18.000 cell populations corresponding to single gene deletion strains in the haploid yeast deletion collection. We report that the fitness of gene knockout near-perfectly (*R^2^=*0.91) predicts their adaptation dynamics under arsenic exposure, leaving virtually no role for dedicated evolvability functions in the corresponding proteins. We tracked the adaptation of another >23.000 yeast gene knockout populations to a diverse range of selection pressures and generalised the almost perfect (*R^2^=*0.72 to 0.98) capacity of initial fitness to predict the rate of adaptation. Finally, we reconstruct mutations in the genes *FPS1*, *ASK10,* and *ARR3*, which together account for almost all arsenic adaptation in wildtype cells, in gene deletions covering a broad fitness range. We show that the predictability of arsenic adaptation can be understood almost entirely as a global epistasis phenomenon where excluding arsenic from cells, through these mutations, is more beneficial in cells with low arsenic fitness regardless of what causes the arsenic defects. The lack of genes with a meaningful effect on the adaptation dynamics of clonally reproducing cell populations diminishes the prospects of developing adjuvant drugs aiming to slow antimicrobial and chemotherapy resistance.

## INTRODUCTION

Clonal adaptive evolution and expansion is the ultimate cause of infectious diseases. Fully or partially asexual populations of mostly microbial pathogens adapt to the tissue environment and immune defence of humans, crop plants, and domesticated animals, as well as to measures taken to prevent or treat infections. Bacterial antibiotic resistance alone may cause millions of deaths annually unless new effective countermeasures are taken (Dadgostar, 2019). The adaptive evolution of somatic cells to their tissue environment and to medical treatments is also the root cause of tumour progression and many cases of chemotherapeutic failure (Merlo et al., 2006). More efficient use of existing therapies, development of new treatments, vaccines, and antiseptics, and reduced human and animal exposure to infection and cancer-causing agents can diminish some of the above threats to human health (S. B. Levy & Marshall, 2004). However, the adaptive evolution of pathogenic, cancerous, and drug-resistant variants is challenging to avoid (Ragheb et al., 2019). Complementary approaches that seek to constrain the capacity of such cell populations to generate and transmit beneficial variation upon which selection can act, i.e., their evolvability, have therefore attracted increasing interest (Payne & Wagner, 2019). These approaches see gene products promoting evolvability as potential chemical targets that can be inhibited to slow cancer or microbial pathogen adaptation to their surrounding human tissue environments and to the medical measures taken against them. The clinical use of such adjuvant drugs could enhance the efficacy of antimicrobial and anticancer treatment regimes, and extend their usefulness by delaying resistance development in patients and microbial populations at large (Ragheb et al., 2019).

The mechanisms underlying evolvability have been explored theoretically (Alberch, 1991; Conrad, 1990; Houle, 1992; Pigliucci, 2008; A. Wagner, 1996; G. P. Wagner & Altenberg, 1996), and experimental studies have demonstrated the involvement of cellular processes such as DNA replication and repair that control the mutation rate (Lynch et al., 2016). Mutator lineages with elevated mutation rates are also often observed in asexually evolving cell populations (Lynch et al., 2016; Sniegowski et al., 1997; Wielgoss et al., 2013), where the linkage of mutator mutations to beneficial alleles they give rise to allow the former to hitchhike to high frequencies. However, while high mutation rates can shorten the time for a selectable beneficial variant to emerge, this occurs at the cost of a higher genetic load of deleterious mutations. Whether elevated mutation rates enhance adaptation rates in natural and clinical contexts is therefore rarely clear and will likely vary with the type and strength of selection pressure (Ram & Hadany, 2012). Alleles promoting larger cell populations, for example, by channelling limited resources into more, rather than larger, cells are expected to lead to the generation of more new variants. So are alleles reducing the generation time (Olson-Manning et al., 2012). Sexually reproducing yeast populations have been shown to adapt faster than asexual by joining beneficial alleles arising in different genomes into one and by releasing such alleles from linked deleterious variants (McDonald et al., 2016). Gene products affecting mating, outbreeding and the recombination rate can therefore be thought of as speeding up adaptation.

Imperfections in transcription, translation and protein degradation (Blake et al., 2003; Elowitz et al., 2002) lead to macromolecular diversity in cell populations that can manifest as heterogeneity in fitness. Even if resulting variants are not transmitted across generations, such diversity may enhance evolvability and speed adaptation. Yeast expresses the self-propagating prion form [*PSI+*] of the translational termination protein Sup35 (True & Lindquist, 2000) when faced with certain environmental challenges (Tyedmers et al., 2008). Self-aggregation of [*PSI+*] reduces the translational termination fidelity, leading to increased rates of stop-codon read-through (Baudin-Baillieu et al., 2014; Eaglestone et al., 1999; Firoozan et al., 1991; Lancaster et al., 2010). The resulting protein variants are cryptic because they reach high abundance only in [*PSI+*] cells, benefiting cell survival and reproduction in adverse conditions (Zabinsky et al., 2019). Simulations support that the [psi-]/[*PSI+*] system can serve as a phenotypic switch that act as an evolutionary capacitor, providing robustness in benign conditions by canalising certain favourable phenotypes while also de-canalising these phenotypes during adverse conditions, when variation is needed (Lancaster et al., 2010; True & Lindquist, 2000). The yeast chaperone Hsp90 canalise phenotypes by chaperoning mutated peptides into a standard protein fold, thereby preventing their expression as phenotypes on the cellular and higher levels (Rutherford & Lindquist, 1998). However, during stress, the higher number of misfolded proteins can overwhelm the Hsp90 system, allowing mutated peptides to be expressed as incorrectly folded proteins and de-canalising the phenotype. This unmasking of beneficial protein variants enhances evolvability, while exposing deleterious protein variants facilitates purging of deleterious mutations from the population (Jarosz & Lindquist, 2010). The chaperone GroEL in *E. coli* and *HSP-110* in *C. elegans* may canalise and de-canalise genetic variation in much the same way (Koneru et al., 2021; Tokuriki & Tawfik, 2009).

Simulations have suggested that gene networks can mask much phenotypic variation. This variation can then be uncovered when critical genes in the network are compromised, or switched off, thereby accelerating adaptation (Bergman & Siegal, 2003). Around 300 yeast genes have been shown to control morphological cell-cell variation, supporting that they have secondary functions as phenotypic stabilisers (Levy & Siegal, 2008). And through their control of diversity in expression, chromatin regulators may be particularly important for generating phenotypic variation (Tirosh et al., 2010).

While it is broadly accepted that certain gene products can promote the generation of variation, whether these variants influence adaptation dynamics is highly unclear. To a large extent, this is because accurately tracking the adaptation dynamics of many cell populations in parallel to screen for genetic effects on adaptation, is challenging. Here we capitalise on a recently introduced high throughput platform for accurate measuring of microbial growth (Zackrisson et al., 2016) to track the cell doubling time adaptation of 18.432 haploid cell populations, corresponding to yeast gene knockouts, to arsenic. We show that the adaptation dynamics of 4639 single gene deletion strains could be near-perfectly predicted by their initial fitness. We generalise this conclusion by evolving a large subset (330 strains) of the deletion collection in four other environments, again finding near-perfect predictability. Finally, we reconstructed mutations in *FP*S*1*, *ASK10* (alternative name: *RGC2*) and *ARR3* (alternative name: *ACR3*) that together account for almost all acquired arsenic resistance in evolving wildtype populations in many deletion strain backgrounds. We show that the slower arsenic adaptation of fitter deletion strains can be explained by a lower effect of these key beneficial mutations. Our results show that gene products affect clonal adaptation of haploid yeast populations almost exclusively through their direct effects on fitness, leaving virtually no room for dedicated evolvability functions.

## RESULTS

### Tracking the adaptation dynamics across >18.000 yeast gene knockout populations

To probe genetic control over adaptation dynamics, we first established an Artificial Laboratory Evolution (ALE) framework to track the adaptation of 18.432 yeast populations in parallel (Fig 1A). In our ALE platform, we expanded populations from ∼50.000 to 2-4 million cells as colonies growing on a nutrient-complete synthetic agar medium, and sampled colonies robotically after three days, after detectable growth had ended. We deposited cell samples on freshly made plates, repeated the batch cultivation, and then cycled each population over 19 rounds of expansion and sampling, corresponding to 80-100 cell generations (Fig 1A). We maintained colonies in a stable environment in bench-top scanners and estimated the population density change in each colony based on measurements of the transmitted light at 20-minute intervals (Zackrisson et al., 2016). We then derived the adaptation for each population as the change in cell doubling time over cell population doublings, the later used as proxy for consecutive cell generations (Fig 1A). Finally, we fitted a Locally Estimated Scatterplot Smoothing (LOESS) regression to the adaptation data for each population to account for technical and environmental variation. This allowed us to estimate the adaptation achieved after each generation for each population (Fig 1A). To survey the effects of individual genes on adaptation dynamics, we evolved the collection of the single yeast gene knockouts (Giaever et al., 2002) clonally in the presence of arsenic in the form of trivalent arsenite (As[III]; 3 mM). Arsenite exposure at this concentration increased the cell doubling time ∼60% (from 2.27 to 3.61 h), of the mean deletion strain (Fig S1). Arsenite is a ubiquitous selection pressure with intracellular toxicity that is countered by a dedicated cellular defence system (Wysocki & Tamás, 2010, 2011). Arsenite enters yeast cells primarily through the Fps1 aquaglyceroporin (Wysocki et al., 2001), whose activity is regulated by Ask10 (Beese et al., 2009; Lee et al., 2013), and is exported primarily by the H^+^ antiporter Arr3 (Wysocki et al., 1997) (Fig 1C). Natural yeast variation in As[III] resistance is explained almost exclusively by translocations, and segmental duplications encompassing the *ARR3* locus (Yue et al., 2017) and lab strain yeast populations exposed to high As[III] adapt either by *ARR3* amplification or by point mutations inactivating Fps1 or Ask10 (Gjuvsland et al., 2016). This reinstates arsenite homeostasis by excluding As[III] from cells and returns the cell doubling time to near pre-stress levels (Gjuvsland et al., 2016). Because these solutions are rapidly encountered even at our relatively moderate population sizes, As[III] adaptation in the lab strain is swift and little afflicted by chance variations, making it an ideal testbed for gene effects on adaptation dynamics. We measured the arsenite adaptation of 4639 yeast populations, corresponding to single gene deletions in the lab strain background that were viable in the presence of 3 mM arsenite. Each gene deletion was represented by 3-6 replicate populations allowing us to account for much of the mutational randomness and measurement error, and we evolved in total 18.432 populations. The 384 wildtype controls achieved 53.6% (cell doubling time) of their final adaptation within the first 25 generations; after that, their adaptation plateaued. They completed only 18.7% of their last adaptation after 75 generations (Fig 1B). This wildtype pattern of adaptation was also shared by the vast majority of faster and slower adapting gene knockouts, and most genes did not measurably affect As[III] adaptation kinetics (Fig 1B). A substantial minority of gene knockouts adapted faster than the wildtype, with improvements distributed along a continuum from marginal to very large (Fig 1B). In contrast, only a few adapted substantially slower than the wildtype adaptation, reflecting that few genes benefitted evolvability appreciably. To reduce the data dimensionality, we focused on the adaptation achieved over shorter (25 generations), medium (50 generations) and longer (75 generations) time spans (Fig 1A). We found the adaptation achieved by gene knockouts at these time points to predict each other well (linear regression coefficient, *R^2^*=0.88 to 0.95). We, therefore, assume that As[III] adaptation across different time spans is dictated by essentially the same biology (Fig 1D, S2), and that conclusions based on the time points above can be generalized.

**Figure 1.**
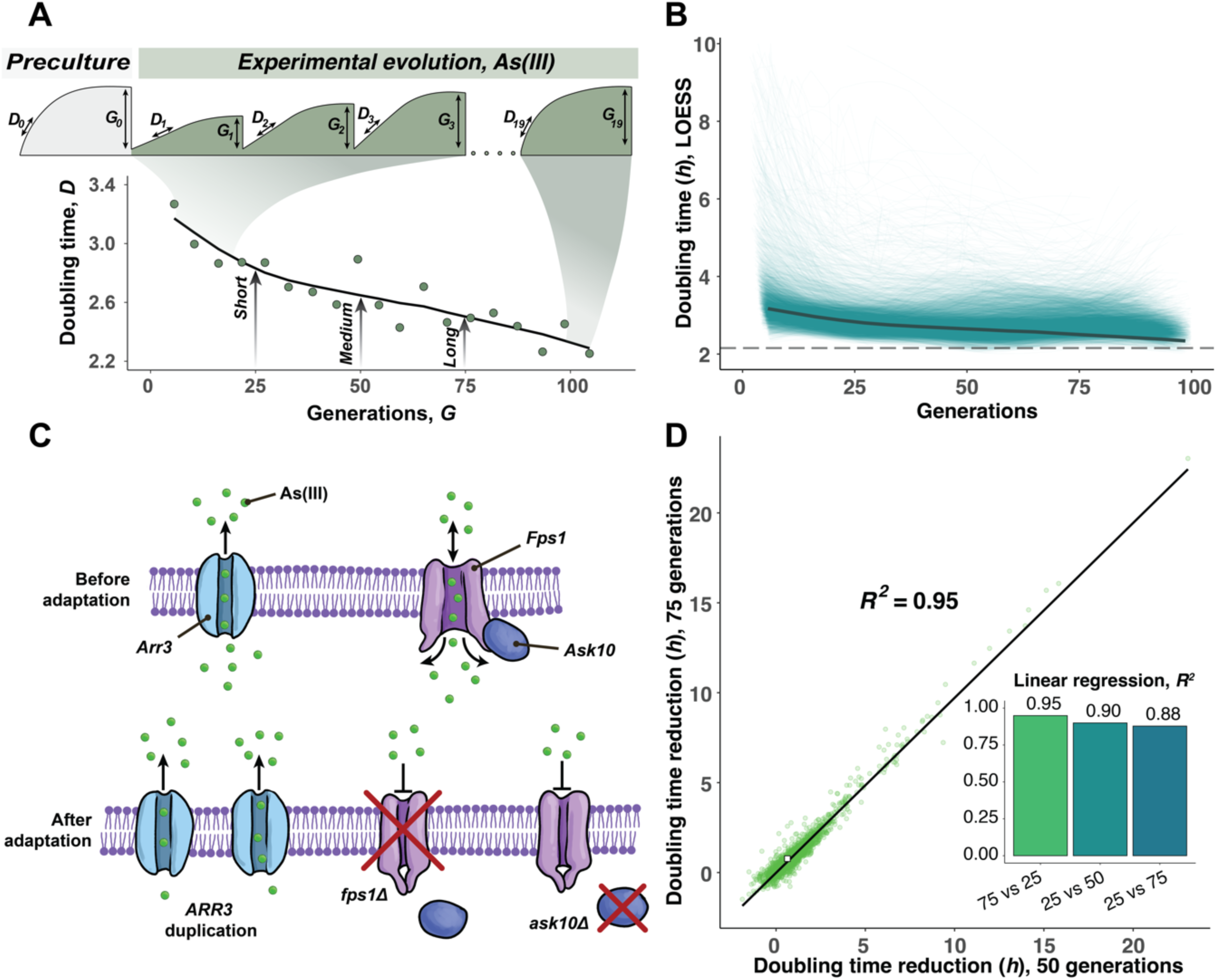
Experimental setup and phenotype extraction. **A)** Experimental evolution and population growth parameters extraction. After an initial preculture cycle on basal medium, populations were transferred to the selective medium. The doubling time (*D*) and generations (*G*) in each cultivation cycle were extracted from the population size growth curve. A LOESS curve was fitted to each growth curve and the adaptation achieved after 25, 50 and 75 generations were extracted from the LOESS fit. The adaptation wildtype colonies (mean values of *n*=384) are shown as an example. **B)** Mean LOESS fitted adaptation curves of all gene deletion mutants, wildtype adaptation is indicated with a thick grey line. The broken line shows the doubling time of wildtype in basal media without stress**. C)** Schematic illustration of arsenite influx and efflux (top) and the three adaptive solutions to exclude arsenite from entering the cell (bottom). **D**) Comparing the doubling time reductions achieved by 4639 yeast deletion strains exposed to arsenite (3mM) over 75 (*y*-axis) and 50 generations (*x*-axis). Mean values (*n*=3-6) are shown, wildtype indicated with a white square (*n*=384). The linear regression and the squared coefficient of linear regression are shown. *Inset:* squared linear regression coefficients from comparing doubling time reductions achieved over 25, 50, and 75 generations of arsenite adaptation.

### Gene knockout fitness near-perfectly predicts arsenite adaptation dynamics

The continuous decline in the adaptation rate for virtually all strains suggested that the fitness of gene knockouts, rather than any evolvability function of the gene, controls the kinetics of adaptation. We probed this conjecture by examining the As[III] adaptation achieved after 25, 50 and 75 generations of all 18.432 cell populations in light of their cell doubling time, as a proxy for fitness, before adaptation. Overall, the cell doubling time before adaptation predicted change in cell doubling time across all evolutionary time spans near perfectly (linear regression coefficient, *R^2^*=0.88-0.91), (Fig 2A, 2B, S3). The prediction accuracy approached or exceeded the repeatability of single replicate measures of adaptation (linear regression coefficient, *R^2^*=0.69-0.72), which is limited only by the measurement error, environmental variation between colony positions, and mutational randomness. Adaptation was dramatically slower for fitter gene knockouts, reflecting a diminishing return of adaptation as fitness improves (Fig 2A, S3). This was not due to fitter gene knockouts reaching a selection limit, dictated, for example, by cell-intrinsic constraints on growth set by ribosome production reaching a maximum, because adapting cell populations still grew slower than unstressed wildtype cell populations (Fig 2A, S3). Some gene knockouts adapted significantly (Students t-test, FDR, *q=*0.05) better (*n=*219-1331) or worse (*n=*3-995) than expected from their arsenite fitness. Still, their deviations from the expectation for those who did were almost uniformly small (median of 0.24-0.30 hours higher and 0.28-0.45 hours lower) and are likely due to unaccounted environmental variation between colony positions rather than intrinsic differences between gene knockouts. Consistent with this assumption, no cellular functions (yeast GO slim, Fisher’s exact test, FDR *q*>0.05) were enriched among these genes. Moreover, genes often suspected of influencing evolvability, such as those encoding DNA repair or protein folding functions, generally adapted as predicted by their fitness (Fig 2A, S3A). This included the Hsp90 chaperone Hsp82/Hsc82, as well as key components in the single-strand break repair (Tdp1), mismatch repair (Msh2), base-excision repair (Mre11), non-homologous end-joining (Yku70), homologous recombination (Rad51) and meiotic recombination (Spo11). We found arsenite adaptation to be largely independent of the initial cell doubling time of gene knockouts in the absence of arsenite (linear regression coefficient, *R^2^=*0.11) and gene knockouts with perfect growth in the absence of arsenite sometimes vastly improved their arsenite growth. This is consistent with the observed arsenite adaptation resulting from improvements in the arsenite cellular biology, rather than from the evolutionary rescue of more general growth defects caused by the gene deletions. Overall, we conclude that the near-perfect predictability of the arsenite adaptation dynamics from the initial arsenite fitness of gene deletion strains leaves virtually no room for separate evolvability functions in the corresponding genes.

**Figure 2.**
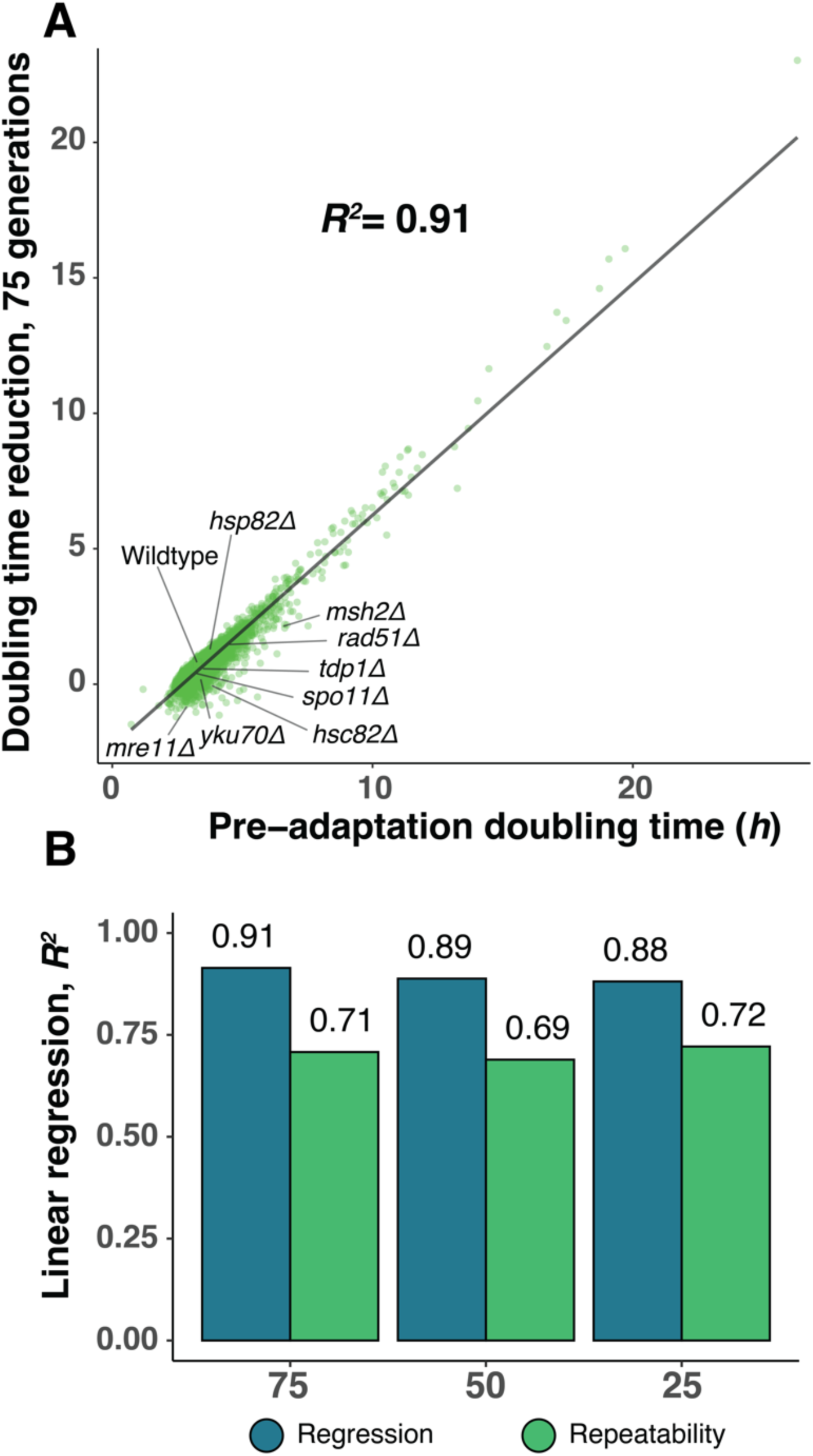
Adaptation of gene deletion strains to arsenite is near perfectly predicted by their fitness. A) The doubling time reduction in 4639 yeast deletion strains exposed to arsenite (3mM) over 75 generations as a function of their pre-adaptation doubling time. Mean values (*n*=3-6) are shown, wildtype indicated with a white square (*n*=384). The linear regression line and the squared coefficient of linear regression are shown. B) Squared coefficient of linear regression comparing the pre-adaptation doubling time and the doubling time reduction over 25, 50 and 75 generations of adaptation to arsenite (blue) and the squared linear regression between replicate measures of doubling time reduction at these time points (green, single replicate repeatability).

### Fitness near-perfectly predicts gene knockout adaptation across a range of selection pressures

The randomness of mutations and the fact that new mutations can be lost due to stochastic genetic drift when still rare in populations likely accounts for some of the observed adaption variation between deletion strains. To reduce this cause of uncertainty, we repeated the arsenite (3 mM As[III]) ALE for 345 deletion strains, covering the complete spectrum of adaptation kinetics, at higher replication (*n=*12-16). The estimates of the adaptation dynamics of these gene knockout populations showed arsenite fitness to virtually perfectly predict all genetic variation in adaptation across the three evolutionary timespans considered (linear regression coefficient, *R^2^=*0.93 to 0.96) (Fig 3A, S4). Some deletion strains deviated in adaptation from that predicted by their initial cell doubling time. Still, their deviations were all small (median of 0.31-0.49 hours higher and 0.63-0.79 hours lower among significant deviations). Next, we asked whether the extraordinary predictive power of fitness on arsenite adaptation was independent of the strength of the arsenite selection. We, therefore, performed ALE on another set of 330 random deletion strains, again at high replication (*n*=12-16) to 4 mM As[III]. The stronger arsenite selection (wildtype cell doubling time increase 6.54 vs 4.86 h at 3 mM) forced five of the slowest gene deletion strains to go extinct. Still, for the remaining 98.5% of gene deletion strains, their initial arsenite fitness again predicted essentially all arsenite genetic variance in adaptation (linear regression coefficient, *R^2^*=0.97-0.98) (Fig 3B, S5). Thus, the outstanding predictive power of fitness on adaptation dynamics persisted also at stronger arsenite selection. Finally, we asked whether fitness predicted the adaptation of gene deletion strains to a similar degree also under selection pressures to which cells adapt through processes other than those that drive arsenite resistance. We therefore repeated the ALE for the second set of 330 random gene knockouts, at high replication (*n*=12-16), under selection imposed by the redox-cycler paraquat (400 mg/L), the immunosuppressant rapamycin (0.25 mg/L), and hyperosmotic stress inducer NaCl (1.25 M). These impair cell doubling time by targeting different aspects of yeast physiology (Table S1), which was underscored by the absence correlation in gene deletion strain growth between environments (pairwise linear regression coefficient, *R^2^<*0.01 to 0.02) (Fig S6). Again, we found the initial cell doubling time of gene deletion strains in the presence of each of these stresses to predict virtually all genetic variation in their subsequent adaptation dynamics, with correlations (linear regression coefficient, *R^2^*=0.72 to 0.98) approaching or exceeding that between replicated measures of adaptation (Fig 3C). Outliers, whose adaptation was imperfectly explained by the initial fitness, were few and their deviations from the predicted adaptation were small. Moreover, cells lacking key DNA repair and protein folding genes adapted as predicted from their initial fitness. Stress adaptation generally occurred at the cost of fitness loss in the absence of the stressor (fig S7), underscoring that the adaptation was not to the background environment but rather to the specific selection agent added. While some cause for caution remains in that this screen did not cover the entire gene deletion collection, the random selection of strains tested gives substantial confidence to the assessment that gene products with dedicated evolvability functions is rare in yeast.

**Figure 3.**
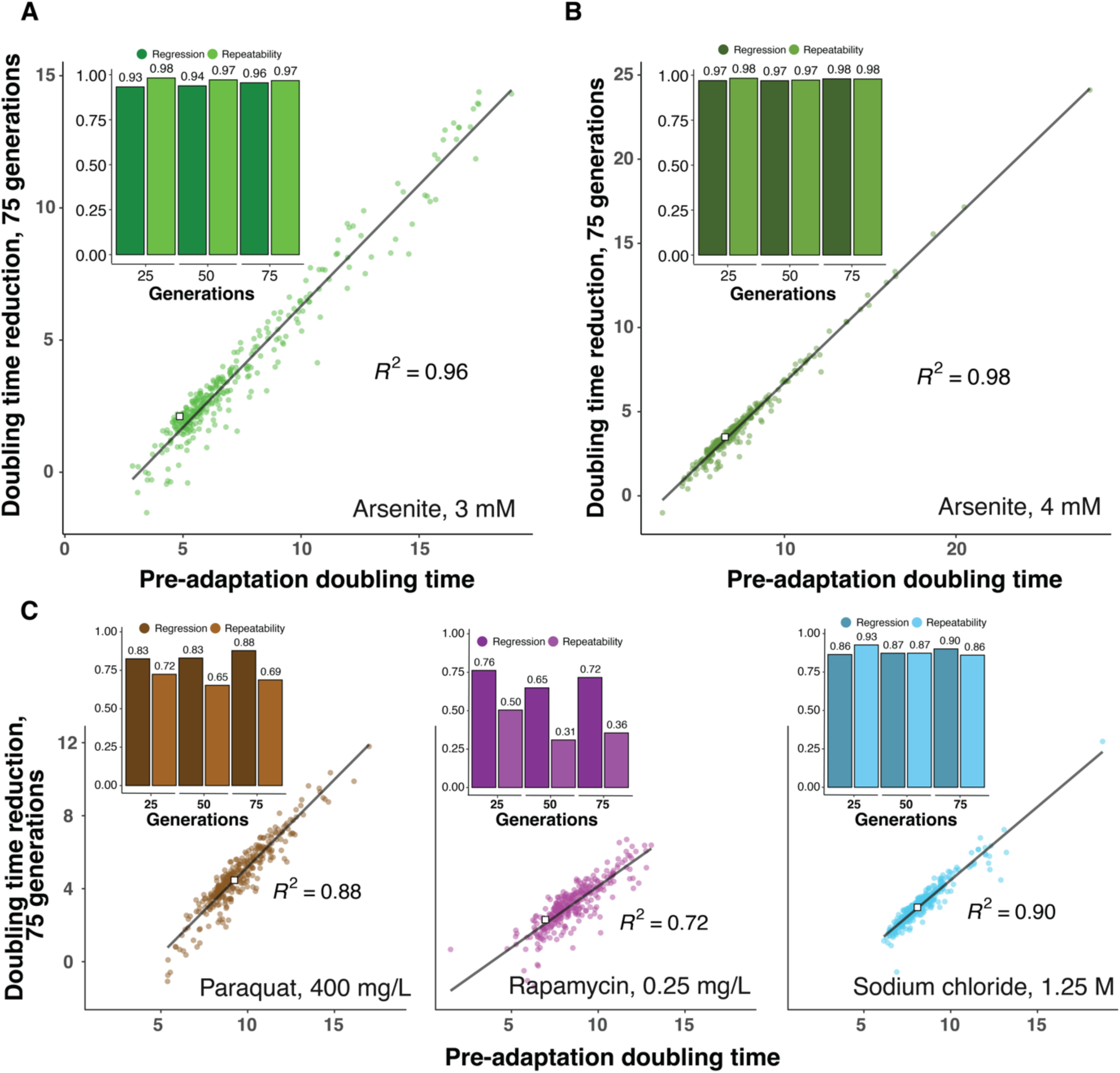
Deletion strain adaptation across a range of selection pressures. Doubling time reductions achieved by 330-345 yeast deletion strains (mean of *n*=12-16) over 75 generations of adaptation to A) arsenite 3 mM, B) arsenite 4 mM and C) paraquat 400 mg/L, rapamycin, 0.25 mg/L and sodium chloride, 1.25 M. White squares indicate the adapting wildtype. Linear regression lines and squared linear regression coefficients are shown. *Inset:* Squared linear regression coefficients comparing pre-adaptation doubling time and the doubling time reduction over 25, 50 and 75 generations of adaptation to the stress indicated (dark) and the squared linear regression between replicate measures of doubling time reduction at these time points (light, single replicate repeatability).

### Diminishing return epistasis dictates yeast arsenite adaptation

Adaptation is expected to decline with increasing fitness, if strongly positive mutations are few, rapidly fixate and become depleted (Barton, 1998; Orr, 1998, 1999). However, recent studies have provided strong support for an alternative explanation for adaptation slowing with increasing fitness: many beneficial mutations are less beneficial in fitter backgrounds (Chou et al., 2011; Couce & Tenaillon, 2015; Khan et al., 2011; Miller, 2019). To test to what extent the latter hypothesis can explain the slower arsenite adaptation, we reconstructed the three mutation types that account for almost all arsenite adaptation in wildtype cells - *ARR3* duplication and *FPS1* and *ASK10* loss, in gene knockout strains covering a broad range of cell doubling times on arsenite. We introduced complete *FPS1* and *ASK10* gene deletions into each of 464 gene knockouts strains by mating, meiosis, and sporulation (Kuzmin et al., 2014) and measured their cell doubling time on 3 mM As[III] in the presence and absence of Fps1 and Ask10. *FPS1* deletion, and in some cases *ASK10* deletion, had a negative impact on cell growth in the absence of arsenite, which varied depending on the genetic background (Fig 4A). This likely reflect the importance of cells being able to export glycerol through the Fps1 channel to maintain osmotic homeostasis (Luyten et al., 1995; Tamas et al., 1999). We accounted for these effects by comparing the doubling time of each strain in the absence and presence of arsenite and then extracting the cell doubling time effect of *FPS1* and *ASK10* deletion, respectively, on this measure of arsenite resistance. Overall, loss of FPS1 conferred greater arsenite resistance than did ASK10 loss (mean of 3.0 vs 1.0 hours, Fig 4B), consistent with the fact that Fps1 regulation involves other proteins besides Ask10 (Ahmadpour et al., 2016; Beese et al., 2009; Lee et al., 2013; Mollapour & Piper, 2007; Thorsen et al., 2006). However, both Fps1 and Ask10 loss conferred much stronger arsenite specific growth benefits to arsenite sensitive than to arsenite resistant gene knockouts. In fact, the increase in arsenite resistance due to either Fps1 or Ask10 loss could be well predicted (linear regression coefficient, *R^2^*=0.52-0.92) by the arsenite fitness of the strain into which the mutations were introduced. For example, removal of Fps1 was highly beneficial in strains lacking the transcription factors Yap1 (regulator of oxidative stress response) and Rpn4 (regulator of proteotoxic stress response) (Rathod et al., 2018), which both are key to cells maintaining fitness on arsenite, while having much smaller effects on strains with unperturbed arsenite homeostasis (Fig 4B-C). We validated that the lesser impact of arsenite adaptive mutations in fitter backgrounds is not a property specific for changes to the Fps1 system by also reconstructing the Arr3 duplication in a subset of the gene deletion strains. We thus introduced an extra *ARR3* gene, carried on a single-copy plasmid, into 140 deletion strains and estimated the beneficial effect of this mutation on arsenite resistance. The pattern of a diminishing return of the *ARR3* duplication in more arsenite resistant deletion strains was also abundantly clear (linear regression coefficient, *R^2^*=0.65) (Fig 4B-C). Overall, the power of the doubling time of deletion strain to predict the arsenite resistance conferred by introducing a Fps1 or Ask10 loss, or Arr3 duplication into this strain was high, again approaching or exceeding the capacity of replicated measures of mutation effects to predict each other (Fig 4C). Thus, diminishing return epistasis well accounted for the variation in the effect size of arsenite beneficial mutations across deletion strains, with unaccounted for variation likely explained by measurement error, environmental variation, or the emergence of random background mutations during the construction process. We conclude that excluding arsenite from cells through *FPS1* or *ASK10* loss of function mutations or through *ARR3* duplication is more beneficial if cells have poor arsenite fitness. Thus, the near-perfect predictive power of fitness on arsenite adaptation dynamics is well explained by diminishing return epistasis.

**Figure 4.**
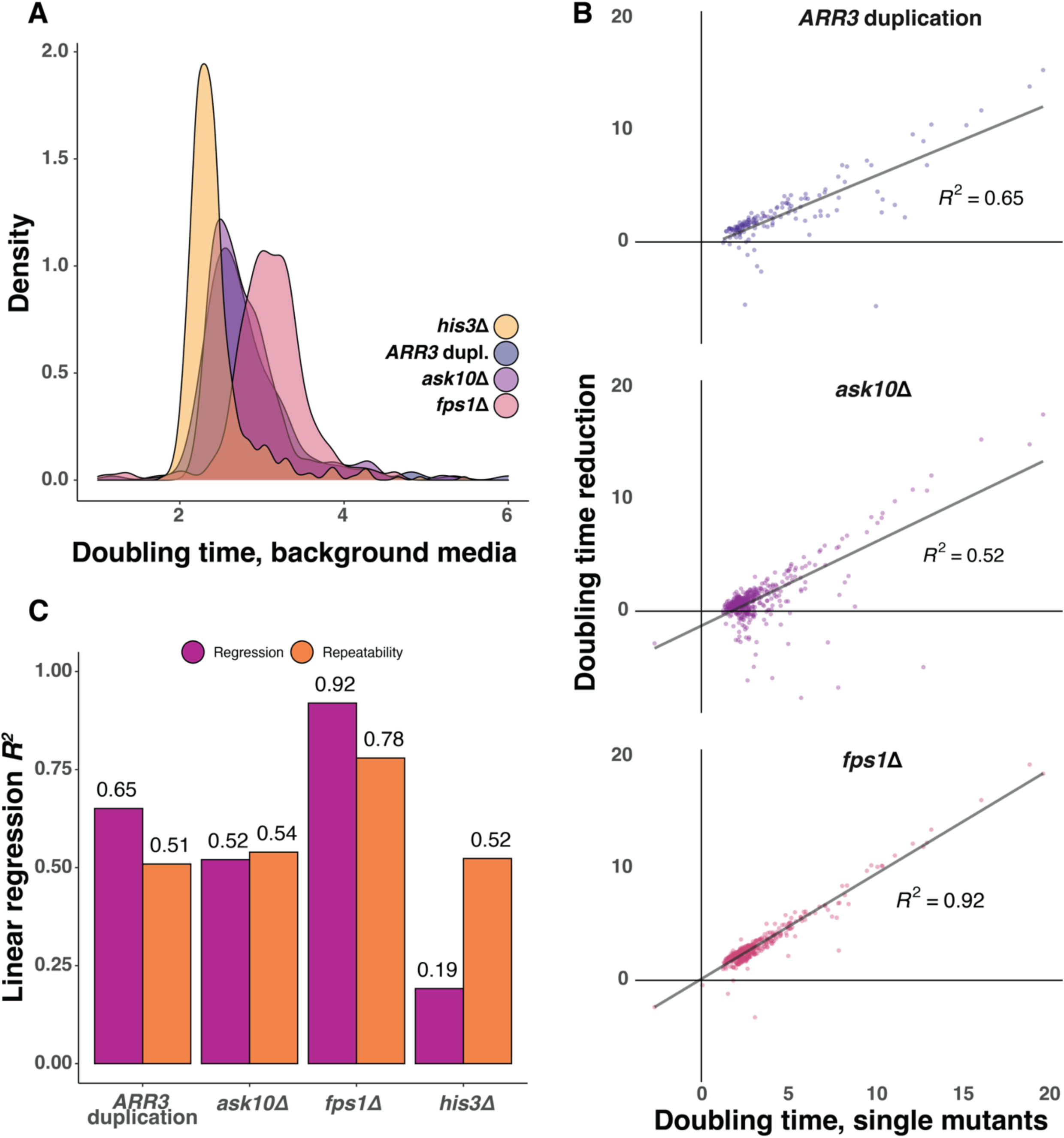
Diminishing return epistasis dictates yeast arsenite adaptation. A) Density distribution of doubling times of gene deletion strain also missing *ASK10* (*n*=468) or *FPS1* (*n*=468) or having an extra *ARR3* copy (*n*=140) in the no stress background media. The same strains also missing the neutral *HIS3* (*n*=468) are shown as a control. Means of *n*=6-612 replicates were used. B) Arsenite (3 mM) specific doubling time reductions when removing *ASK10* or *FPS1* or inserting an extra *ARR3* copy arsenite 3 mM exposure in single gene deletion strains with different doubling times on arsenite (*x*-axis). Linear regressions and the squared regression coefficient are indicated. C) Squared linear regression coefficients between the arsenite (3mM) specific doubling time reductions when removing *ASK10* or *FPS1* or inserting an extra *ARR3* copy arsenite 3 mM exposure in single gene deletion strains and the initial doubling times on arsenite of these single gene deletion strains. The squared linear regression between replicate measures of the former is shown for comparison (single replicate repeatability).

## DISCUSSION

### Evolvability genes have no meaningful role in the clonal adaptation of haploid yeast populations

We here tracked the arsenite adaptation of almost all viable single gene deletions in a yeast lab strain background. We showed that essentially all genetically encoded variation in adaptation dynamics could be explained by their variation in fitness, with less fit gene deletion strains adapting much faster than fitter. Using large subsets of gene deletion strains and high replication, we showed that this conclusion is valid also for stronger selection and for a variety of biologically distinct selection pressures. The general tendency for adaptation to decline with increasing fitness should come as no surprise and has been reported before, in smaller studies on virus, bacteria, and yeast (Couce & Tenaillon, 2015; Jerison et al., 2017; Lukačišinová et al., 2020; MacLean et al., 2010). What is remarkable in this more exhaustive study is the capacity of fitness to account for virtually all heritable variation in the adaptation kinetics: it leaves minimal room for dedicated evolvability functions in gene products to have had a meaningful impact on adaptation rates. This is further underscored by the fact that strains lacking gene products argued to have such roles, such as functions controlling how fast novel genetic variation emerges in cells or penetrates as phenotypic variation, adapted with almost precisely the speed predicted by their fitness. Clearly, there are limits to the extent to which these findings should be generalised. At least in the case of arsenite, our ALE populations operated in a strong selection, strong mutation regime. The abundant access to strongly beneficial arsenite resistance mutations in *FPS1*, *ASK10* and *ARR3* means that these solutions invariably will be found very rapidly. Hence, elevations of the mutation rates, through e.g., removal of DNA replication and repair components will, almost per definition, do little to speed up adaptation (Gjuvsland et al., 2016). In smaller populations, or populations with access to fewer strong mutations, the mutation rate will be a stronger limiting factor on the adaptation rate. Elevations of the former may thus have a greater effect in such population (Sniegowski & Gerrish, 2010). Second, adaptation in haploid cell populations may not perfectly capture the effects of evolvability factors on adaptation rate in diploid cell populations. In diploid cell populations, recessive variants, such as most loss-of-function mutations, can drive clonal adaptation only if first converted into homo- or hemizygotic states (Vázquez-García et al., 2017). Evolvability factors that promote homo- or hemizygosity, such as those inducing gene conversion, chromosome segment deletions or non-reciprocal translocations, may affect adaptation rates in diploid cell populations that we are unable to capture. Likewise, as our populations are asexual, we fail to capture any evolvability effects of mating or inbreeding or of genes increasing meiotic (or mitotic in the case of parasexual fungal pathogens) recombination rates. Finally, transformation or conjugation promoting factors that enhance the horizontal transmission of genes, which at least in bacteria can have large effects on adaptation rates (Alalam et al., 2020; Graf et al., 2019), will be overlooked here. Bearing these caveats in mind, our finding that the >4.600 probed genes possessed no functions with a substantial impact on adaption rate is nevertheless quite remarkable and calls for caution when considering the evolutionary importance of evolvability in general. More specifically, it questions the idea that natural selection has acted extensively on living systems to promote the establishment of dedicated evolvability associated functions. And in terms of human health, it diminishes the prospects of evolvability functions becoming valuable targets for drugs that are given in concert with antimicrobials and chemotherapeutics to slow resistance development.

### Global diminishing return epistasis dictates arsenite adaptation dynamics

We could explain the slower arsenite adaptation in fitter gene deletions by a diminishing return epistasis. Benefits of excluding arsenite from the cell, through loss of function mutations in the Fps1 arsenite importer or its positive regulator Ask10 or duplications of the arsenite efflux protein Arr3, continuously decreased with increasing fitness of the deletion strains in which these mutations were reconstructed. The smaller mutation effect sizes were evident in fit strains lacking a wide variety of gene functions, as well as in wildtype cells. They are therefore not reflections of modular epistasis (Wei & Zhang, 2019), within the arsenite efflux or influx systems, but of a global epistasis where the effect of Fps1 loss, Ask10 loss and Arr3 duplication depend on a broad range of other variants in the genome in which mutations emerge (Kryazhimskiy et al., 2014). Interpreted within the context of Fps1, Ask10 and Arr3 function, such a global epistasis makes perfect biological sense: if arsenite is effectively excluded from cells, through a dramatically reduced influx (Fps1, Ask10) or increased efflux (Arr3), it becomes irrelevant what other variants affecting arsenite homeostasis that present in a genome as the toxicity of arsenite is entirely intracellular (Fig 1C). Other gene products could conceivably affect arsenite uptake, with e.g., exported glutathione that binds extracellular arsenite and prevents its entry (Thorsen et al., 2012), and a small amount of arsenite entering cells through hexose transporters (Liu et al., 2004). But their small effects on arsenite resistance, together with the high rate of loss-of-function mutations in Fps1 and Ask10 and of Arr3 duplications, means that the latter almost invariably will drive arsenite adaptation in populations matching the size of our ALE colonies (Gjuvsland et al., 2016).

Diminishing return epistasis may not always be the sole genetic determinant of adaptation kinetics. Tumours with an inactivated P-glycoprotein drug efflux pump, the normal site for resistance mutations to some chemotherapeutics, adapt slowly to these treatments, reflecting a more specific genetic interaction (Binkhathlan & Lavasanifar, 2013). Similarly, disrupting a drug efflux pump can slow the adaptation of *E. coli* populations exposed to antibiotics by shifting them onto evolutionary paths where some mutations reduce the effect size of key resistance mutations (Lukačišinová et al., 2020). Nevertheless, both theoretical (Kryazhimskiy et al., 2009, 2014; Perfeito et al., 2014; Vaishnav et al., 2022) and smaller-scale studies in bacteria (Chou et al., 2011; Khan et al., 2011; MacLean et al., 2010; Wang et al., 2016), virus (Levy & Siegal, 2008; MacLean et al., 2010; Rokyta et al., 2011), yeast (Kryazhimskiy et al., 2014; Wei & Zhang, 2019) and multicellular fungi (Schoustra et al., 2016) support a strong role of diminishing return epistasis in adaptation. Our findings, precisely tracking the adaptation of a near complete genome-wide deletion collection, underscores that the power and generality of diminishing indeed are immense.

### Concluding remarks

Accurate tracking of adaptation in cell population ultimately rests on the precise counting of cells. However, counting cells at sufficiently high resolution and with sufficiently high accuracy in tens of thousands of evolving cell populations is challenging. This is primarily because light transmission through a cell population, the standard proxy for cell density, does not scale linearly with the population size, and unlike in small-scale experiments, this cannot be solved by continuously diluting populations (Warringer & Blomberg, 2003). This gives rise to large measurement errors for both cell division times and the number generations based, giving rise to substantial confounding effects. Our ALE platform, which relies on the Scan-o-matic system, use built-in calibration functions and local regression to translate the transmitted light to actual population size (Zackrisson et al., 2016). With the caveat that the calibration functions need to be adjusted to account for the light scattering and absorbing properties of the specific cell type, the ALE platform is suitable for a broad range of microorganisms and eco-evolutionary questions. In that sense, it may help usher areas of evolutionary biology that previously have only been amenable to moderate-scale studies into the realm of high throughput experimentation.

## ACKNOWLEDGMENTS

**Funding:** This work was supported by the Swedish Research Council (2015-05427, 2018-03638 and 2018-03453) and the Swedish Research Council for Environment, Agricultural Sciences and Spatial Planning (Formas: grant number 942–2015–376) to M.J.T.

## MATERIALS AND METHODS

### Yeast strains

The haploid BY4741 single gene deletion collection (*MAT**a***; *his3*Δ*1*; *leu2*Δ*0*; *met15*Δ*0*; *ura3*Δ*0*; *GeneX*::*kanMX*) and its parental strain BY4741 (wildtype) were used throughout all adaptive evolution experiments (Giaever et al., 2002). To construct double knockout lines, single knockout strains were crossed with one of several BY4742 query strains (*MATα*; *GeneX*::*natMX4*; *can1*Δ::*STE2pr-Sp_his5*; *lyp1*Δ; *his3*Δ*1*; *leu2*Δ0; *ura3*Δ*0*; *met15*Δ*0*). For gene duplication strains, single knockout lines were transformed with a centromeric plasmid (MoBY-*ARR3*: *YPR201W*) containing the *ARR3* gene (Hei Ho et al., 2013).

### Yeast cultivation conditions

Frozen glycerol stocks of yeast strains were recovered on YPD (Yeast Peptone Dextrose) medium supplemented with G418 (Geneticin, 200 mg/L). Wildtype cells were recovered on YPD without added G418 since the strain does not carry the selectable KanMX cassette. Except from revival of frozen stocks, yeast strains were cultivated on a Synthetic Complete medium (SC) composed of 0.14% Yeast Nitrogen Base (CYN2210, ForMedium), 0.5% NH4SO4, 0.077% Complete Supplement Mixture (CSM; DCS0019, ForMedium), 2.0% (w/v) glucose, pH buffered to 5.8 with 1.0% (w/v) succinic acid and 0.6% (w/v) NaOH and addition of 2.0% (w/v) agar for solid medium. Selective environments consisted of the basic SC media composition, but with addition of one of the following stress components: 3 mM arsenite ([As III]; NaAs2O3), 4 mM arsenite, 0.25 mg/L rapamycin, 400 mg/L paraquat (methylviologen; N,N-dimethyl-4-4′-bipiridinium dichloride) and 1.25 M sodium chloride (NaCl). Agar dissolved in deionised water was sterilised by autoclavation and brought to approximately 60 °C when stock solutions of medium components and stressors were added and mixed to homogeneity. Singer Plus plates were cast on a solid surface by adding 50 mL medium. Plates were allowed to dry at room temperature for two days before use.

All yeast strains were stored at -80 °C in 20% glycerol and cultivated at 30 °C. Yeast populations were subsampled and transferred to fresh plates by robotic pinning (ROTOR HDA, Singer Instruments, UK).

### Strain construction

#### Double gene deletions

Double deletion strains were constructed using the Synthetic Genetic Array (SGA) method (Kuzmin et al., 2014). Query strains, deleted for one of the following: *FPS1*, *ASK10*, *URA3*, *HO* and *HIS3*, were prepared as lawns by spreading 800 µl liquid culture on YPD agar, supplemented with adenine (120 mg/L) and clonNAT (100 mg/L). Single deletion strains were robotically pinned in 384 array formats on separate YPD agar plates supplemented with G418 (200 mg/L). Plates were incubated at 30 °C for two days, ensuring sufficient growth. The query strain lawns were transferring cells from lawns in 384 formats to fresh YPD media (in the absence of selective agents). The single deletion strain array was transferred on top of these, mixed robotically, and incubated for one day at 22 °C. To select for diploids, the resulting *MAT***a**/α diploid zygotes were transferred to YPD agar supplemented with G418 (200 mg/L) and clonNAT (100 mg/L) and incubated for two days at 30 °C. The resulting diploid array was transferred to enriched sporulation agar plates (Kuzmin et al., 2014) and incubated at 22 °C for 14 days to ensure high sporulation efficiency. To select for *MAT***a** meiotic haploid progeny, the spores were transferred to SC agar without His/Arg/Lys and without succinate buffer, but supplemented with canavanine (50 mg/L) and thialysine (100 mg/L) and with monosodium glutamic ac (MSG, 1 g/L) instead of ammonium sulfate. The array was incubated for two days at 30 °C. The spores were then transferred to SC MSG without His/Arg/Lys but supplemented with canavanine/thialysine/G418 (concentrations as above) and incubated for two days at 30 °C. In the final selection step, the arrays were transferred to SC MSG without His/Arg/Lys but supplemented with canavanine/thialysine/G418/clonNAT (concentrations as above) and incubated for two days at 30 °C. To ensure that the resulting array consisted of haploid double mutants, the entire array was transferred a second time to fresh plates containing the same media.

#### Gene duplications

Strains were constructed in a microtiter plate using the standard Lithium acetat (LiAc)/single stranded carrier DNA/polyethylene glycol (PEG) method (Gietz & Schiestl, 2007). Strains were cultivated for three days on solid media in 96 array format. A transformation mix was prepared for each 96 well plate consisting of 1 M LiAc, 1.5 mL; single stranded carrier DNA (2 g/L, denatured at 95 °C for 5 min, then transferred to ice), 2 mL; plasmid (≥100 ng/well) dissolved in sterile deionised water, 1.5 mL. For each transformation, 50 µL transformation mix was added per well. Yeast cells were transferred robotically to the 96 well plate and mixed into suspension. To each transformation reaction, 100 µL PEG 3350 (50% w/v) were added and mixed. The plates were incubated at 42 °C for 1.5 h. To recover the transformants, plates were centrifuged at 1500 *g* for 10 min and the supernatant discarded. The pelleted cells were suspended by pipetting in 50 µL liquid selective media (SC-URA), 20 µL of the cell suspension were transferred to a fresh 96 well microtiter plate containing 150 µL selective media (SC-URA) and incubated at 30 °C for three days. Transformants were stored in 20% glycerol at -80 °C.

### Artificial laboratory evolution procedure

Deletion mutants in 384 arrays were recovered on YPD + G418 media from -80 °C glycerol stocks. After three days of cultivation at 30 °C, strains were subsampled and replicated three times on preculture plates, leaving every fourth position empty, 384 positions per plate in total. Wildtype spatial controls were similarly prepared on YPD media. Adaptation was performed without adding wild type spatial controls, leaving every fourth position empty. In preparation for phenotyping, the adapting deletion strains were subsampled and, together with wildtype controls in the fourth position transferred to preculture plates containing the stressor. After three days of cultivation at 30 °C, precultures were transferred to fresh plates and transferred to scanners, and growth phenotypes were acquired.

Along with every round of adaptation, a new preculture was made, and fresh wildtype spatial controls were added. The wildtype controls were added using 384 pads, while the adapting deletion strain populations were added with 1536 pads. This caused a more extensive initial population size favouring the wildtype spatial controls on preculture plates, preventing usage of these plates directly for phenotyping. Transferral from preculture plates to were performed using 1536 format pads. Fresh plates were kept at room temperature, and plates pinned for phenotyping were transferred to scanners within 20 min of transferral of yeast populations.

### Measuring population doubling time

We tracked the growth for all cell populations using the Scan-o-matic system (Zackrisson et al., 2016) version 2.2 (https://github.com/Scan-o-Matic/scanomatic/releases/tag/v2.2). Plates were maintained undisturbed without lids for 72 h in high-definition desktop scanners (Epson Perfection V800 PHOTO scanners, Epson Corporation, UK), placed inside dark, humid, and temperature-controlled (30 °C) thermostatic cabinets. With four plates placed in each scanner, images were acquired using SANE (Scanner Access Now Easy) by transmissive scanning at 600 dpi. The plates were held in position by an acrylic glass fixture. Pixel intensity was normalised and standardised across the different scanners and experiments using a transmissive grayscale calibration strip (LaserSoft IT8 Calibration Target, LaserSoft Imaging, Germany).

The pixel intensity of the greyscale calibration strips was compared to the manufacturer’s values; this allowed normalisation of variations in the light intensity of the transmission scan. Colonies were detected by the software using a virtual grid across each plate, with intersections matching the centre of each colony. At the intersections, colonies and surrounding areas were segmented to determine the local background and pixel intensities. The pixel intensity was converted to total cell numbers using a pre-defined, independent calibration function, based on both spectroscopic and flow cytometer measurements. From this calibration, population size growth curves were obtained. The series of population size measurements were smoothed in a two-step procedure to remove random noise variation. First, local spikes in each curve were removed by a median filter. Second, the remaining local noise was reduced by a Gaussian filter. The steepest slope in each growth curve was identified using a local regression over five consecutive time points, converting the slopes into population size doubling times. Growth curves with poor quality were automatically detected and manually inspected before exclusion. We estimated the number of cell generations passed in each growth cycle as the total number of population doublings, between the last and the first population size estimates. Population parameters were extracted as numerical values from all growth curves that passed the quality requirements. We fitted a locally estimated scatterplot smoothing (LOESS) regression to the adaptation data for each adapting population to account for technical and environmental variation, allowing estimation of the adaptation achieved at each stage of evolution for each population.

### Resource availability

The authors declare that all data supporting the findings of this study are available within the paper as Supplemental Information Data S1-S7, which can be previewed at DOI: 10.17632/r5kz3kj6f2.1.

### Materials availability

All stored unique strains and stored populations generated in this study are available from the Lead Contact without restriction.

## Supplemental dataset descriptions

**Data S1:** reports dose-response growth data for gene deletion strains exposed to different degrees of arsenite stress. **Data S2:** reports growth data, before LOESS fitting, for the complete collection of BY4741 gene deletion strains during adaptive evolution in 3 mM of arsenite. **Data S3:** reports growth data, after LOESS fitting, for the complete collection of BY4741 gene deletion strains after 25, 50 and 75 generations of adaptive evolution in 3 mM of arsenite. **Data S4:** reports growth data, before LOESS fitting, of BY4741 gene deletion strains during adaptive evolution across a range of selection pressures. **Data S5:** reports growth data, after LOESS fitting, of BY4741 gene deletion strains after 25, 50 and 75 generations across a range of selection pressures. **Data S6:** reports growth data, of BY4741 gene deletion strains on background media before and after adaptation to a range of selection pressures. **Data S7:** reports growth data, of single and double mutation strains in Synthetic complete medium, and arsenite, 3 mM.

---

**Figure S1.**
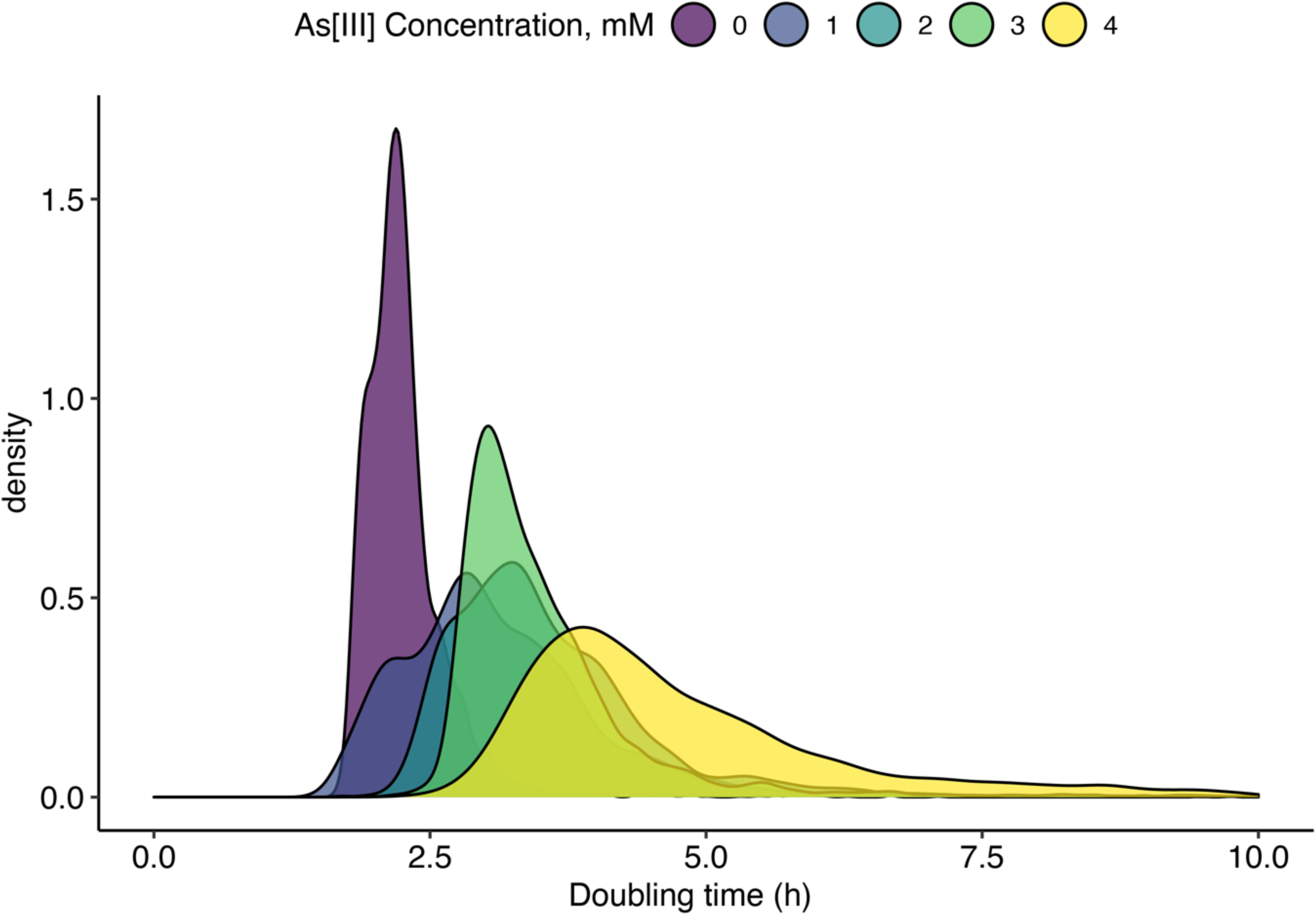
Doubling times of single gene deletion strains exposed to arsenite. Density distribution of mean doubling time of all single gene deletion strains exposed to different concentrations of arsenite (mean of *n*=3-6).

**Figure S2.**
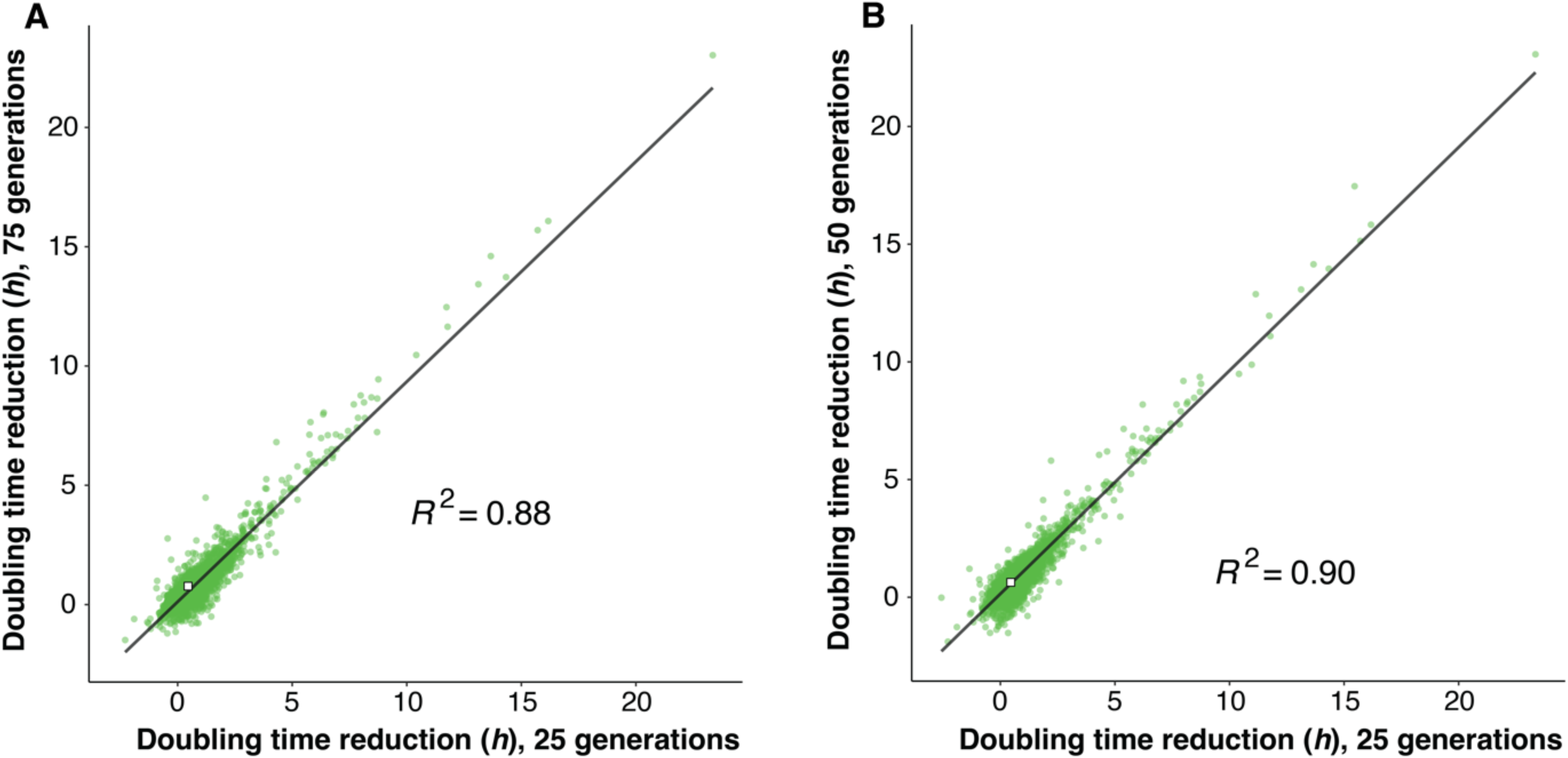
Arsenite adaptation of yeast deletion strain at different time points of adaptation. Comparing the doubling time reductions achieved by 4639 yeast deletion strains (mean of *n*=3-6) exposed to arsenite (3mM) A) over 75 (y-axis) vs. 25 generations (x-axis) and B) over 50 (y-axis) vs 25 generations (x-axis). The linear regression and the squared coefficient of linear regression (also shown in Fig. 1D) are shown.

**Figure S3.**
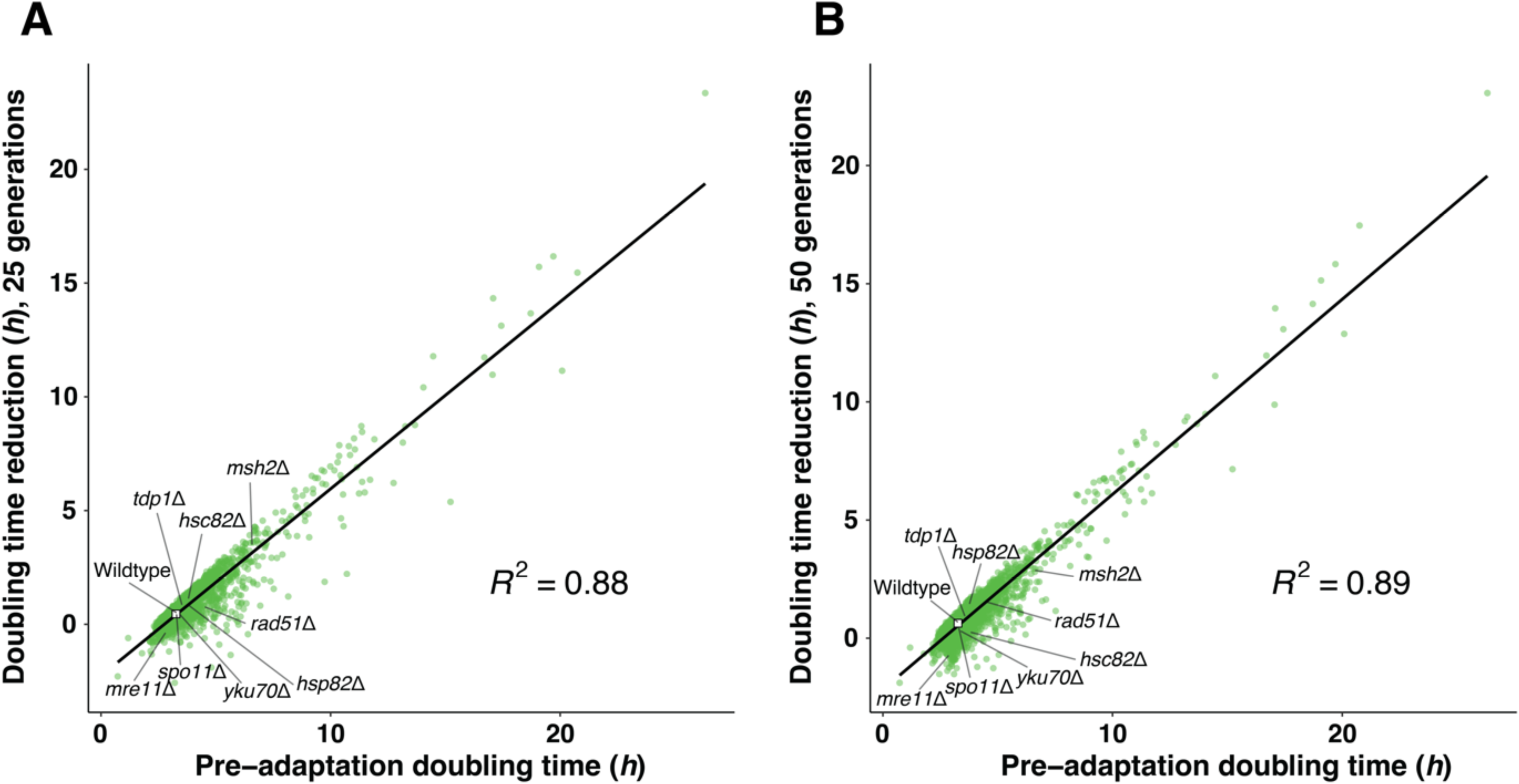
Adaptation of gene deletion strains to arsenite is near perfectly predicted by their fitness. The doubling time reduction in 4639 yeast deletion strains exposed to arsenite (3mM) over A) 25 generations and B) 50 generations, as a function of their pre-adaptation doubling time. Mean values (*n*=3-6) are shown, wildtype indicated with a white square (*n*=384). The linear regression line and the squared coefficient of linear regression (also shown in Fig 2A) are shown.

**Figure S4.**
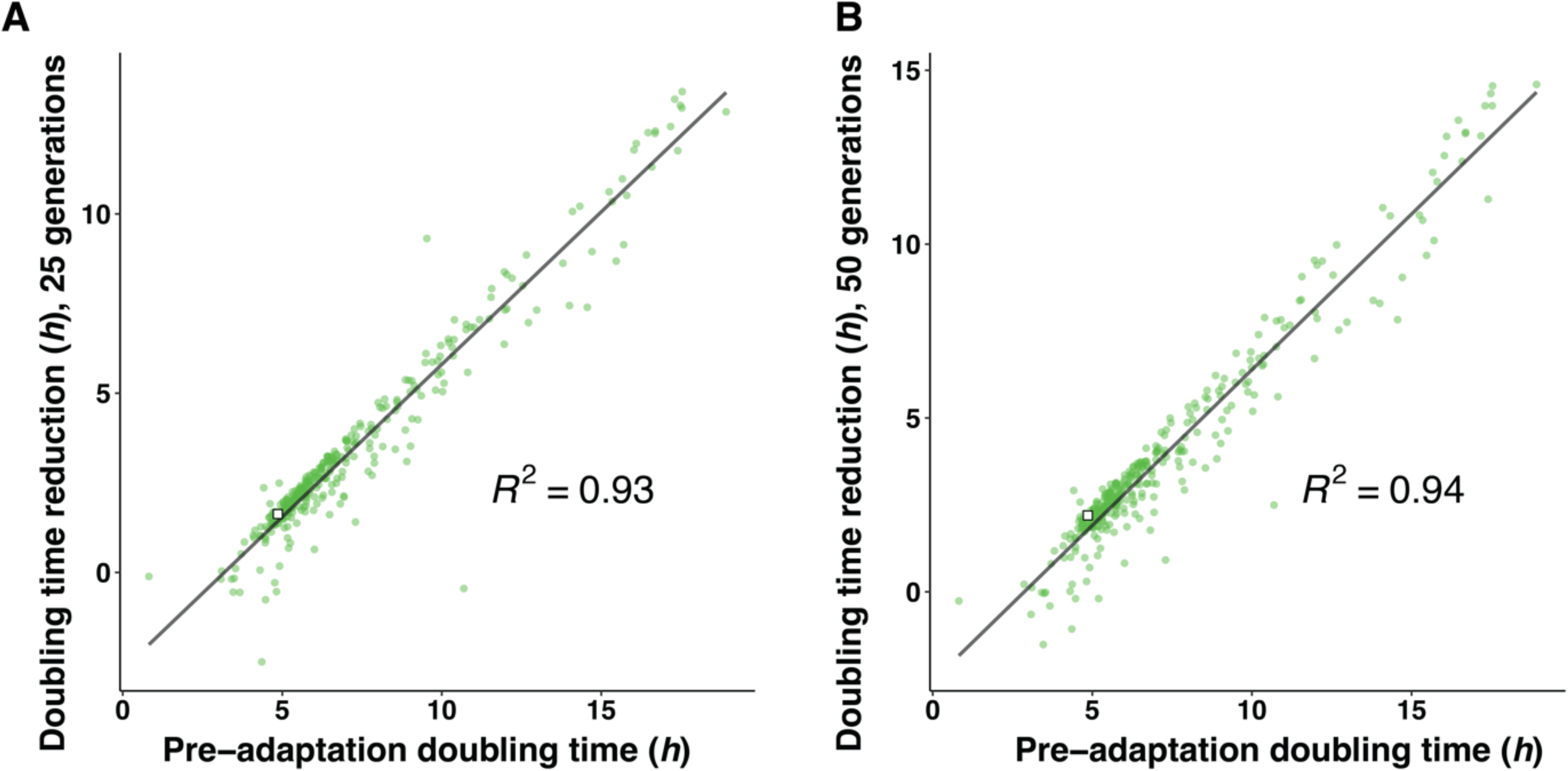
Adaptation of yeast deletion strains to 3 mM As[III] at high replication is near perfectly predicted by initial fitness. Doubling time reductions achieved by 345 yeast deletion strains over A) 25 and B) 50 generations of adaptation to arsenite (3 mM), as a function of their pre-adaptation doubling time. Mean values of *n*=12 replicates are shown. White squares indicate the adapting wildtype (mean of *n*=468). Linear regression lines and squared linear regression coefficients (also shown in Fig 3A) are shown.

**Figure S5.**
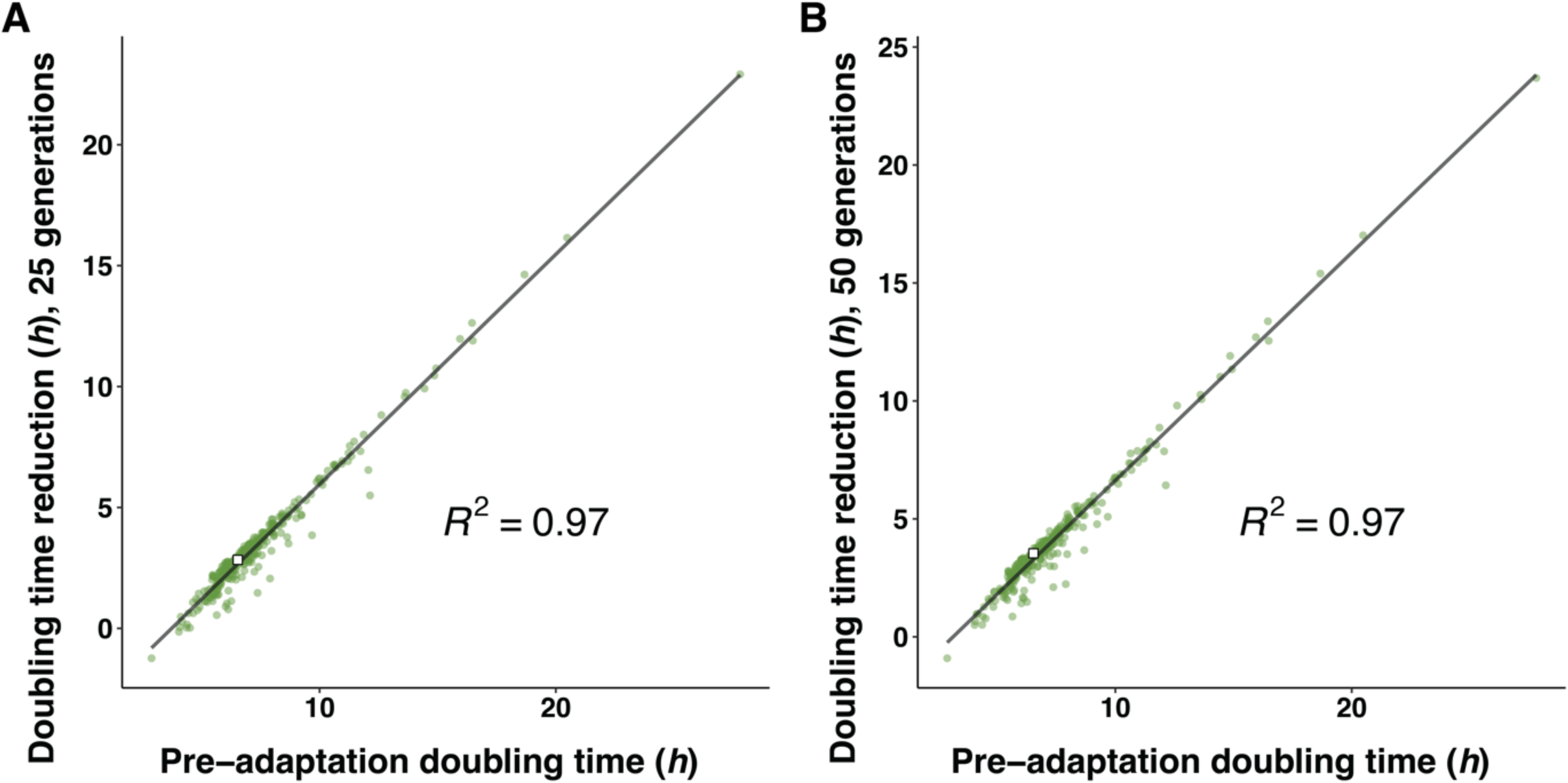
Adaptation of yeast deletion strains to 4 mM As[III] at high replication is near perfectly predicted by initial fitness. Doubling time reductions achieved by 330 yeast deletion strains over A) 25 and B) 50 generations of adaptation to arsenite (4 mM), as a function of their pre-adaptation doubling time. Mean values of *n*=12 replicates are shown. White squares indicate the adapting wildtype (mean of *n*=648). Linear regression lines and squared linear regression coefficients (also shown in Fig 3B) are shown.

**Figure S6.**
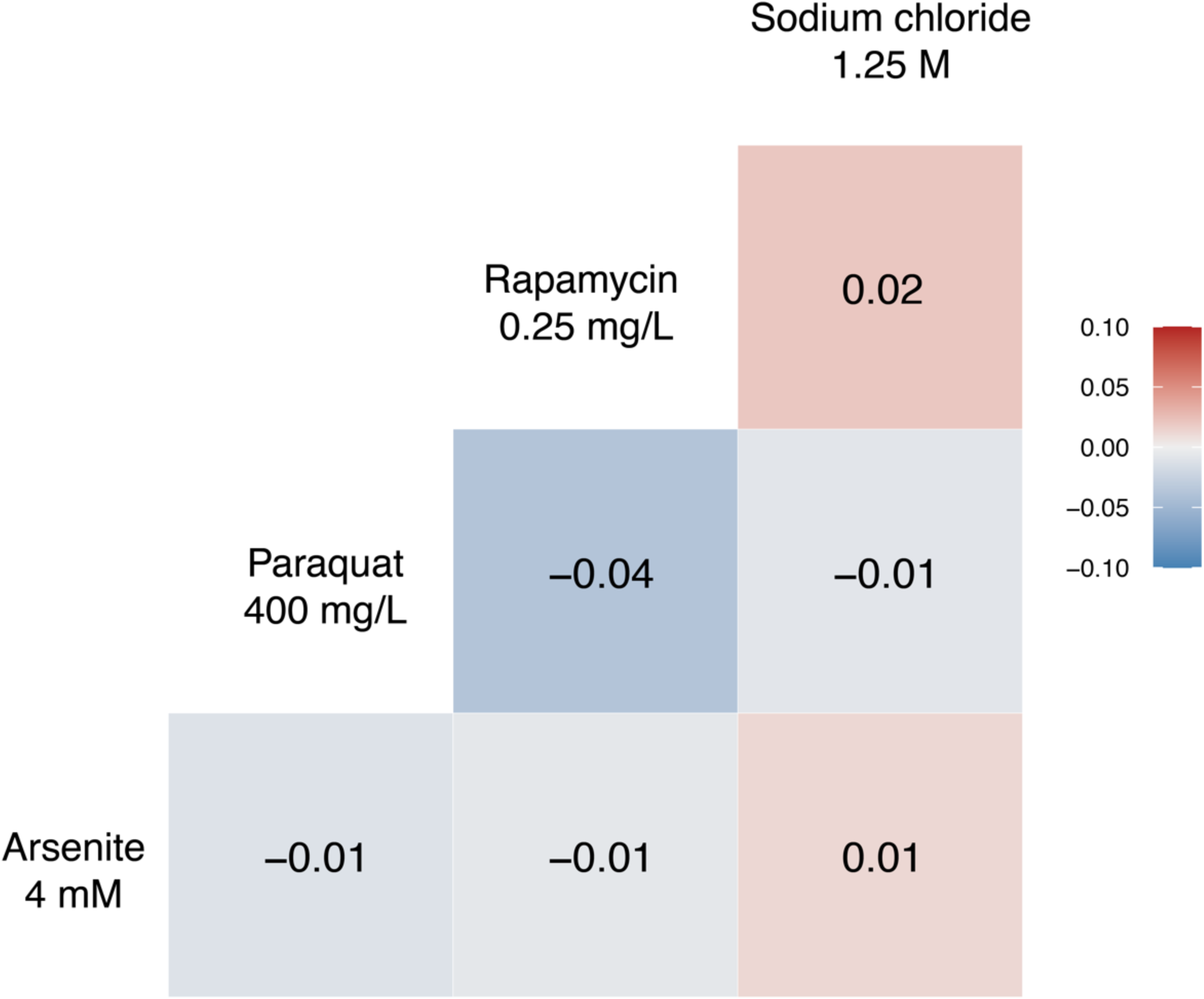
Yeast deletion strain adaptations to different selection pressures are uncorrelated. We compared the doubling time reductions achieved by 330 yeast deletion strains (mean of *n*=12) over 75 generations of adaptation to arsenite 4 mM, paraquat 400 mg/L, rapamycin, 0.25 mg/L and sodium chloride, 1.25 M. Numbers indicate Pearson’s correlation coefficient, *r*, for each pairwise comparison of environments.

**Figure S7.**
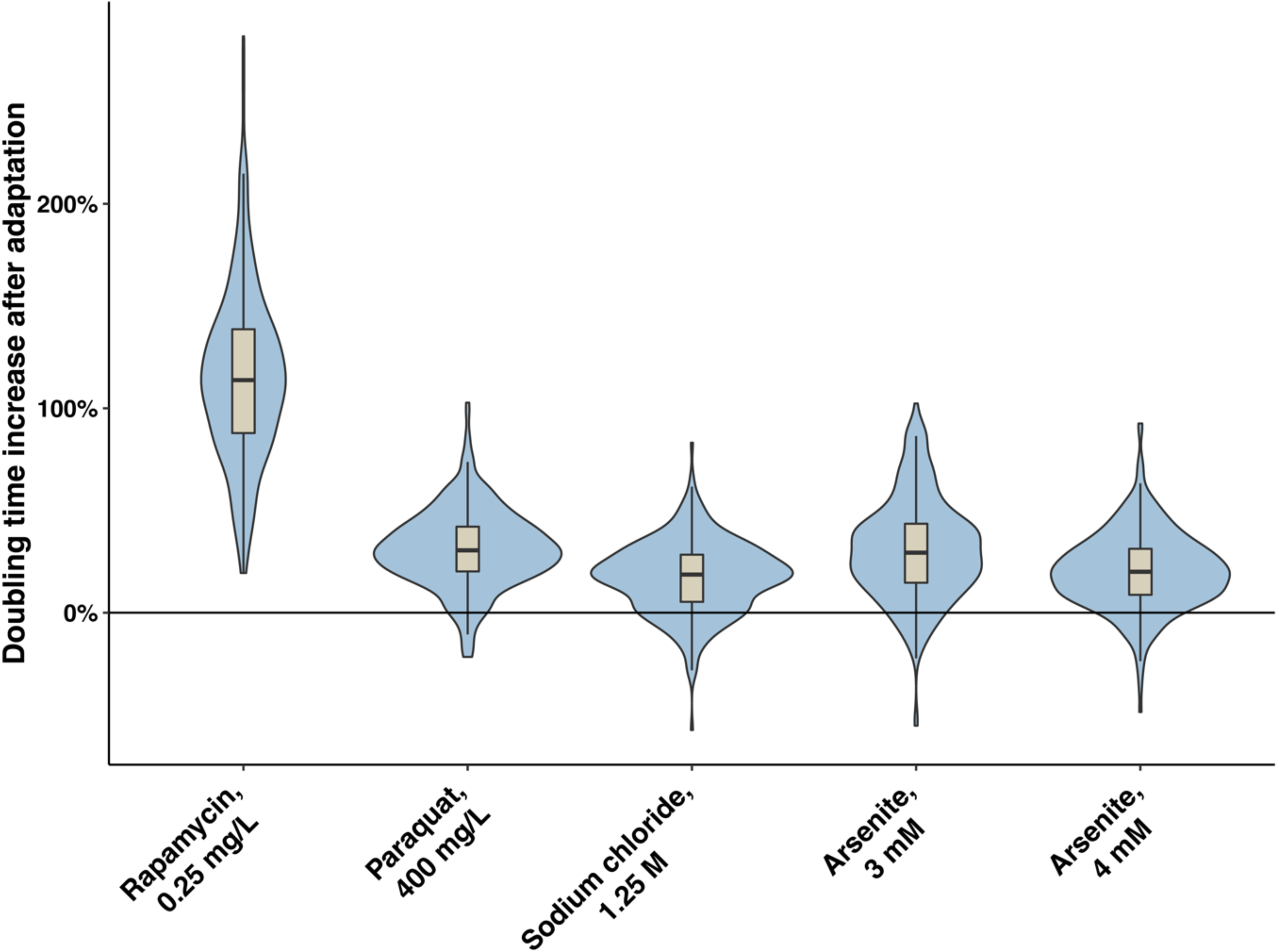
Adaptation of yeast deletion strains to stress incurs a cost in absence of stress. Violin-box diagrams shows how much (in %) the doubling time in absence of stress (SC background medium) increases in yeast deletion strains (*n*=330-345) adapted over 75 generations to a range of selection pressures. Mean values (*n*=12-16) were used. The grey box shows the 25^th^ to 75^th^ quantile. Whiskers shows values within the 1.5 interquartile range from the hinge. Horizontal line indicates the median. Outline shows the probability distribution.

**Table S1:** List of selection agents used in the evolution experiments

| Selection | Compound | Concentration(s) | Type |
| --- | --- | --- | --- |
| Arsenite (III) stress | Sodium arsenite<br>(NaAsO<sub>2</sub>) | 3 and 4 mM | Toxic metalloid |
| Mitochondrial<br>superoxide stress | Paraquat | 400 mg/L | Mitochondrial<br>superoxide producer |
| Rapamycin stress | Sirolimus | 0.25 mg/L | TOR-inhibitor |
| Osmotic stress | Sodium chloride (NaCl) | 1.25 M | Salt |

## Notes

https://doi.org/10.17632/r5kz3kj6f2.1

### Competing Interest Statement

The authors have declared no competing interest.

## REFERENCES

1. Ahmadpour, D., Maciaszczyk-Dziubinska, E., Babazadeh, R., Dahal, S., Migocka, M., Andersson, M., Wysocki, R., Tamás, M. J., & Hohmann, S. (2016). The mitogen-activated protein kinase Slt2 modulates arsenite transport through the aquaglyceroporin Fps1. FEBS Letters, 590(20), 3649–3659. https://doi.org/10.1002/1873-3468.12390

2. Alalam, H., Graf, F. E., Palm, M., Abadikhah, M., Zackrisson, M., Boström, J., Fransson, A., Hadjineophytou, C., Persson, L., Stenberg, S., Mattsson, M., Ghiaci, P., Sunnerhagen, P., Warringer, J., & Farewell, A. (2020). A High-Throughput Method for Screening for Genes Controlling Bacterial Conjugation of Antibiotic Resistance. MSystems, 5(6), 1– 14. https://doi.org/10.1128/mSystems.01226-20

3. Alberch, P. (1991). From genes to phenotype: dynamical systems and evolvability. Genetica, 84(1), 5–11. https://doi.org/10.1007/BF00123979

4. Barton, N. (1998). The geometry of adaptation. Nature, 395(6704), 751–752. https://doi.org/10.1038/27338

5. Baudin-Baillieu, A., Legendre, R., Kuchly, C., Hatin, I., Demais, S., Mestdagh, C., Gautheret, D., & Namy, O. (2014). Genome-wide Translational Changes Induced by the Prion [PSI+]. Cell Reports, 8(2), 439–448. https://doi.org/10.1016/j.celrep.2014.06.036

6. Beese, S. E., Negishi, T., & Levin, D. E. (2009). Identification of Positive Regulators of the Yeast Fps1 Glycerol Channel. PLoS Genetics, 5(11), e1000738. https://doi.org/10.1371/journal.pgen.1000738

7. Bergman, A., & Siegal, M. L. (2003). Evolutionary capacitance as a general feature of complex gene networks. Nature, 424(6948), 549–552. https://doi.org/10.1038/nature01765

8. Binkhathlan, Z., & Lavasanifar, A. (2013). P-glycoprotein Inhibition as a Therapeutic Approach for Overcoming Multidrug Resistance in Cancer: Current Status and Future Perspectives. Current Cancer Drug Targets, 13(3), 326–346. https://doi.org/10.2174/15680096113139990076

9. Blake, W. J., KÆrn, M., Cantor, C. R., & Collins, J. J. (2003). Noise in eukaryotic gene expression. Nature, 422(6932), 633–637. https://doi.org/10.1038/nature01546

10. Chou, H.-H., Chiu, H.-C., Delaney, N. F., Segre, D., & Marx, C. J. (2011). Diminishing Returns Epistasis Among Beneficial Mutations Decelerates Adaptation. Science, 332(6034), 1190–1192. https://doi.org/10.1126/science.1203799

11. Conrad, M. (1990). The geometry of evolution. Biosystems, 24(1), 61–81. https://doi.org/10.1016/0303-2647(90)90030-5

12. Couce, A., & Tenaillon, O. A. (2015). The rule of declining adaptability in microbial evolution experiments. Frontiers in Genetics, 6(MAR), 1–7. https://doi.org/10.3389/fgene.2015.00099

13. Dadgostar, P. (2019). Antimicrobial resistance: implications and costs. Infection and Drug Resistance, 12, 3903–3910. https://doi.org/10.2147/IDR.S234610

14. Eaglestone, S. S., Cox, B. S., & Tuite, M. F. (1999). Translation termination efficiency can be regulated in Saccharomyces cerevisiae by environmental stress through a prion-mediated mechanism. The EMBO Journal, 18(7), 1974–1981. https://doi.org/10.1093/emboj/18.7.1974

15. Elowitz, M. B., Levine, A. J., Siggia, E. D., & Swain, P. S. (2002). Stochastic gene expression in a single cell. *Science (New York*, N.Y.), 297(5584), 1183–1186. https://doi.org/10.1126/science.1070919

16. Firoozan, M., Grant, C. M., Duarte, J. A. B., & Tuite, M. F. (1991). Quantitation of readthrough of termination codons in yeast using a novel gene fusion assay. Yeast, 7(2), 173–183. https://doi.org/10.1002/yea.320070211

17. Giaever, G., Chu, A. M., Ni, L., Connelly, C., Riles, L., Véronneau, S., Dow, S., Lucau-Danila, A., Anderson, K., André, B., Arkin, A. P., Astromoff, A., el Bakkoury, M., Bangham, R., Benito, R., Brachat, S., Campanaro, S., Curtiss, M., Davis, K., … Johnston, M. (2002). Functional profiling of the Saccharomyces cerevisiae genome. Nature, 418(6896), 387–391. https://doi.org/10.1038/nature00935

18. Gietz, R. D., & Schiestl, R. H. (2007). High-efficiency yeast transformation using the LiAc/SS carrier DNA/PEG method. Nature Protocols, 2(1), 31–34. https://doi.org/10.1038/nprot.2007.13

19. Gjuvsland, A. B., Zörgö, E., Samy, J. K., Stenberg, S., Demirsoy, I. H., Roque, F., Maciaszczyk-Dziubinska, E., Migocka, M., Alonso-Perez, E., Zackrisson, M., Wysocki, R., Tamás, M. J., Jonassen, I., Omholt, S. W., & Warringer, J. (2016). Disentangling genetic and epigenetic determinants of ultrafast adaptation. Molecular Systems Biology, 12(12), 892. https://doi.org/10.15252/msb.20166951

20. Graf, F. E., Palm, M., Warringer, J., & Farewell, A. (2019). Inhibiting conjugation as a tool in the fight against antibiotic resistance. Drug Development Research, 80(1), 19–23. https://doi.org/10.1002/ddr.21457

21. Hei Ho, C., Magtanong, L., Barker, S. L., Gresham, D., Nishimura, S., Natarajan, P., Koh, J. L. Y., Porter, J., Gray, C. A., Andersen, R. J., Giaever, G., Nislow, C., Andrews, B., Botstein, D., Graham, T. R., Yoshida, M., & Boone, C. (2013). A molecular barcoded yeast ORF library enables mode-of-action analysis of bioactive compounds. Nat Biotechnol, 18(9), 1199–1216. https://doi.org/10.1016/j.micinf.2011.07.011.Innate

22. Houle, D. (1992). Comparing evolvability and variability of quantitative traits. Genetics, 130(1), 195–204. https://doi.org/10.1093/genetics/130.1.195

23. Jarosz, D. F., & Lindquist, S. (2010). Hsp90 and Environmental Stress Transform the Adaptive Value of Natural Genetic Variation. Science, 330(6012), 1820–1824. https://doi.org/10.1126/science.1195487

24. Jerison, E. R., Kryazhimskiy, S., Mitchell, J. K., Bloom, J. S., Kruglyak, L., & Desai, M. M. (2017). Genetic variation in adaptability and pleiotropy in budding yeast. ELife, 6. https://doi.org/10.7554/eLife.27167

25. Khan, A. I., Dinh, D. M., Schneider, D., Lenski, R. E., & Cooper, T. F. (2011). Negative Epistasis Between Beneficial Mutations in an Evolving Bacterial Population. Science, 332(6034), 1193–1196. https://doi.org/10.1126/science.1203801

26. Koneru, S. L., Hintze, M., Katsanos, D., & Barkoulas, M. (2021). Cryptic genetic variation in a heat shock protein modifies the outcome of a mutation affecting epidermal stem cell development in C. elegans. Nature Communications, 12(1), 3263. https://doi.org/10.1038/s41467-021-23567-1

27. Kryazhimskiy, S., Rice, D. P., Jerison, E. R., & Desai, M. M. (2014). Global epistasis makes adaptation predictable despite sequence-level stochasticity. Science, 344(6191), 1519– 1522. https://doi.org/10.1126/science.1250939

28. Kryazhimskiy, S., Tkǎcik, G., & Plotkin, J. B. (2009). The dynamics of adaptation on correlated fitness landscapes. Proceedings of the National Academy of Sciences of the United States of America, 106(44), 18638–18643. https://doi.org/10.1073/pnas.0905497106

29. Kuzmin, E., Sharifpoor, S., Baryshnikova, A., Costanzo, M., Myers, C. L., Andrews, B. J., & Boone, C. (2014). Synthetic Genetic Array Analysis for Global Mapping of Genetic Networks in Yeast. In J. S. Smith & D. J. Burke (Eds.), Yeast Genetics: Methods and Protocols (pp. 143–168). Springer New York. https://doi.org/10.1007/978-1-4939-1363-3_10

30. Lancaster, A. K., Bardill, J. P., True, H. L., & Masel, J. (2010). The Spontaneous Appearance Rate of the Yeast Prion [PSI +] and Its Implications for the Evolution of the Evolvability Properties of the [PSI +] System. Genetics, 184(2), 393–400. https://doi.org/10.1534/genetics.109.110213

31. Lee, J., Reiter, W., Dohnal, I., Gregori, C., Beese-Sims, S., Kuchler, K., Ammerer, G., & Levin, D. E. (2013). MAPK Hog1 closes the S. cerevisiae glycerol channel Fps1 by phosphorylating and displacing its positive regulators. Genes & Development, 27(23), 2590–2601. https://doi.org/10.1101/gad.229310.113

32. Levy, S. B., & Marshall, B. (2004). Antibacterial resistance worldwide: causes, challenges and responses. Nature Medicine, 10(12 Suppl), S122–9. https://doi.org/10.1038/nm1145

33. Levy, S. F., & Siegal, M. L. (2008). Network Hubs Buffer Environmental Variation in Saccharomyces cerevisiae. PLoS Biology, 6(11), e264. https://doi.org/10.1371/journal.pbio.0060264

34. Liu, Z., Boles, E., & Rosen, B. P. (2004). Arsenic Trioxide Uptake by Hexose Permeases in Saccharomyces cerevisiae. Journal of Biological Chemistry, 279(17), 17312–17318. https://doi.org/10.1074/jbc.M314006200

35. Lukačišinová, M., Fernando, B., & Bollenbach, T. (2020). Highly parallel lab evolution reveals that epistasis can curb the evolution of antibiotic resistance. Nature Communications, 11(1), 1–14. https://doi.org/10.1038/s41467-020-16932-z

36. Luyten, K., Albertyn, J., Skibbe, W. F., Prior, B. A., Ramos, J., Thevelein, J. M., & Hohmann, S. (1995). Fps1, a yeast member of the MIP family of channel proteins, is a facilitator for glycerol uptake and efflux and is inactive under osmotic stress. The EMBO Journal, 14(7), 1360–1371. https://doi.org/10.1002/j.1460-2075.1995.tb07122.x

37. Lynch, M., Ackerman, M. S., Gout, J. F., Long, H., Sung, W., Thomas, W. K., & Foster, P. L. (2016). Genetic drift, selection and the evolution of the mutation rate. Nature Reviews Genetics, 17(11). https://doi.org/10.1038/nrg.2016.104

38. MacLean, R. C., Perron, G. G., & Gardner, A. (2010). Diminishing returns from beneficial mutations and pervasive epistasis shape the fitness landscape for rifampicin resistance in Pseudomonas aeruginosa. Genetics, 186(4), 1345–1354. https://doi.org/10.1534/genetics.110.123083

39. McDonald, M. J., Rice, D. P., & Desai, M. M. (2016). Sex speeds adaptation by altering the dynamics of molecular evolution. Nature, 531(7593), 233–236. https://doi.org/10.1038/nature17143

40. Merlo, L. M. F., Pepper, J. W., Reid, B. J., & Maley, C. C. (2006). Cancer as an evolutionary and ecological process. Nature Reviews. Cancer, 6(12), 924–935. https://doi.org/10.1038/nrc2013

41. Miller, C. R. (2019). The treacheries of adaptation. Science, 366(6464), 418–419. https://doi.org/10.1126/science.aaz5189

42. Mollapour, M., & Piper, P. W. (2007). Hog1 Mitogen-Activated Protein Kinase Phosphorylation Targets the Yeast Fps1 Aquaglyceroporin for Endocytosis, Thereby Rendering Cells Resistant to Acetic Acid. Molecular and Cellular Biology, 27(18), 6446–6456. https://doi.org/10.1128/MCB.02205-06

43. Olson-Manning, C. F., Wagner, M. R., & Mitchell-Olds, T. (2012). Adaptive evolution: Evaluating empirical support for theoretical predictions. Nature Reviews Genetics, 13(12), 867–877. https://doi.org/10.1038/nrg3322

44. Orr, H. A. (1998). The Population Genetics of Adaptation: The Distribution of Factors Fixed during Adaptive Evolution. Evolution, 52(4), 935. https://doi.org/10.2307/2411226

45. Orr, H. A. (1999). The evolutionary genetics of adaptation: a simulation study. Genetical Research, 74(3), 207–214. https://doi.org/10.1017/s0016672399004164

46. Payne, J. L., & Wagner, A. (2019). The causes of evolvability and their evolution. Nature Reviews Genetics, 20(1), 24–38. https://doi.org/10.1038/s41576-018-0069-z

47. Perfeito, L., Sousa, A., Bataillon, T., & Gordo, I. (2014). Rates of fitness decline and rebound suggest pervasive epistasis. Evolution, 68(1), 150–162. https://doi.org/10.1111/evo.12234

48. Pigliucci, M. (2008). Is evolvability evolvable? Nature Reviews Genetics, 9(1), 75–82. https://doi.org/10.1038/nrg2278

49. Ragheb, M. N., Thomason, M. K., Hsu, C., Nugent, P., Gage, J., Samadpour, A. N., Kariisa, A., Merrikh, C. N., Miller, S. I., Sherman, D. R., & Merrikh, H. (2019). Inhibiting the Evolution of Antibiotic Resistance. Molecular Cell, 73(1), 157–165.e5. https://doi.org/10.1016/j.molcel.2018.10.015

50. Ram, Y., & Hadany, L. (2012). The evolution of stress-induced hypermutation in asexual populations. Evolution, 66(7), 2315–2328. https://doi.org/10.1111/j.1558-5646.2012.01576.x

51. Rathod, J., Tu, H.-P., Chang, Y.-I., Chu, Y.-H., Tseng, Y.-Y., Jean, J.-S., & Wu, W.-S. (2018). YARG: A repository for arsenic-related genes in yeast. PloS One, 13(7), e0201204. https://doi.org/10.1371/journal.pone.0201204

52. Rokyta, D. R., Joyce, P., Caudle, S. B., Miller, C., Beisel, C. J., & Wichman, H. A. (2011). Epistasis between Beneficial Mutations and the Phenotype-to-Fitness Map for a ssDNA Virus. PLoS Genetics, 7(6), e1002075. https://doi.org/10.1371/journal.pgen.1002075

53. Rutherford, S. L., & Lindquist, S. (1998). Hsp90 as a capacitor for morphological evolution. Nature, 396(6709), 336–342. https://doi.org/10.1038/24550

54. Schoustra, S., Hwang, S., Krug, J., & de Visser, J. A. G. M. (2016). Diminishing-returns epistasis among random beneficial mutations in a multicellular fungus. Proceedings of the Royal Society B: Biological Sciences, 283(1837), 20161376. https://doi.org/10.1098/rspb.2016.1376

55. Sniegowski, P. D., & Gerrish, P. J. (2010). Beneficial mutations and the dynamics of adaptation in asexual populations. Philosophical Transactions of the Royal Society B: Biological Sciences, 365(1544), 1255–1263. https://doi.org/10.1098/rstb.2009.0290

56. Sniegowski, P. D., Gerrish, P. J., & Lenski, R. E. (1997). Evolution of high mutation rates in experimental populations of E. coli. Nature, 387(6634), 703–705. https://doi.org/10.1038/42701

57. Tamas, M. J., Luyten, K., Sutherland, F. C. W., Hernandez, A., Albertyn, J., Valadi, H., Li, H., Prior, B. A., Kilian, S. G., Ramos, J., Gustafsson, L., Thevelein, J. M., & Hohmann, S. (1999). Fps1p controls the accumulation and release of the compatible solute glycerol in yeast osmoregulation. Molecular Microbiology, 31(4), 1087–1104. https://doi.org/10.1046/j.1365-2958.1999.01248.x

58. Thorsen, M., Di, Y., Tängemo, C., Morillas, M., Ahmadpour, D., van der Does, C., Wagner, A., Johansson, E., Boman, J., Posas, F., Wysocki, R., & Tamás, M. J. (2006). The MAPK Hog1p Modulates Fps1p-dependent Arsenite Uptake and Tolerance in Yeast. Molecular Biology of the Cell, 17(10), 4400–4410. https://doi.org/10.1091/mbc.e06-04-0315

59. Thorsen, M., Jacobson, T., Vooijs, R., Navarrete, C., Bliek, T., Schat, H., & Tamás, M. J. (2012). Glutathione serves an extracellular defence function to decrease arsenite accumulation and toxicity in yeast. Molecular Microbiology, 84(6), 1177–1188. https://doi.org/10.1111/j.1365-2958.2012.08085.x

60. Tirosh, I., Reikhav, S., Sigal, N., Assia, Y., & Barkai, N. (2010). Chromatin regulators as capacitors of interspecies variations in gene expression. Molecular Systems Biology, 6(435), 435. https://doi.org/10.1038/msb.2010.84

61. Tokuriki, N., & Tawfik, D. S. (2009). Chaperonin overexpression promotes genetic variation and enzyme evolution. Nature, 459(7247), 668–673. https://doi.org/10.1038/nature08009

62. True, H. L., & Lindquist, S. L. (2000). A yeast prion provides a mechanism for genetic variation and phenotypic diversity. Nature, 407(6803), 477–483. https://doi.org/10.1038/35035005

63. Tyedmers, J., Madariaga, M. L., & Lindquist, S. (2008). Prion Switching in Response to Environmental Stress. PLoS Biology, 6(11), e294. https://doi.org/10.1371/journal.pbio.0060294

64. Vaishnav, E. D., de Boer, C. G., Molinet, J., Yassour, M., Fan, L., Adiconis, X., Thompson, D. A., Levin, J. Z., Cubillos, F. A., & Regev, A. (2022). The evolution, evolvability and engineering of gene regulatory DNA. Nature, 603(March). https://doi.org/10.1038/s41586-022-04506-6

65. Vázquez-García, I., Salinas, F., Li, J., Fischer, A., Barré, B., Hallin, J., Bergström, A., Alonso-Perez, E., Warringer, J., Mustonen, V., & Liti, G. (2017). Clonal Heterogeneity Influences the Fate of New Adaptive Mutations. Cell Reports, 21(3), 732–744. https://doi.org/10.1016/j.celrep.2017.09.046

66. Wagner, A. (1996). Does evolutionary plasticity evolve? Evolution, 50(3), 1008–1023. https://doi.org/10.1111/j.1558-5646.1996.tb02342.x

67. Wagner, G. P., & Altenberg, L. (1996). Complex adaptations and the evolution of evolvability. Evolution, 50(3), 967–976. https://doi.org/10.1111/j.1558-5646.1996.tb02339.x

68. Wang, Y., Diaz Arenas, C., Stoebel, D. M., Flynn, K., Knapp, E., Dillon, M. M., Wünsche, A., Hatcher, P. J., Moore, F. B. G., Cooper, V. S., & Cooper, T. F. (2016). Benefit of transferred mutations is better predicted by the fitness of recipients than by their ecological or genetic relatedness. Proceedings of the National Academy of Sciences, 113(18), 5047–5052. https://doi.org/10.1073/pnas.1524988113

69. Warringer, J., & Blomberg, A. (2003). Automated screening in environmental arrays allows analysis of quantitative phenotypic profiles in Saccharomyces cerevisiae. Yeast, 20(1), 53–67. https://doi.org/10.1002/yea.931

70. Wei, X., & Zhang, J. (2019). Patterns and Mechanisms of Diminishing Returns from Beneficial Mutations. Molecular Biology and Evolution, 36(5), 1008–1021. https://doi.org/10.1093/molbev/msz035

71. Wielgoss, S., Barrick, J. E., Tenaillon, O., Wiser, M. J., Dittmar, W. J., Cruveiller, S., Chane-Woon-Ming, B., Médigue, C., Lenski, R. E., & Schneider, D. (2013). Mutation rate dynamics in a bacterial population reflect tension between adaptation and genetic load. Proceedings of the National Academy of Sciences of the United States of America, 110(1), 222–227. https://doi.org/10.1073/pnas.1219574110

72. Wysocki, R., Bobrowicz, P., & Ułaszewski, S. (1997). The Saccharomyces cerevisiae ACR3 Gene Encodes a Putative Membrane Protein Involved in Arsenite Transport. Journal of Biological Chemistry, 272(48), 30061–30066. https://doi.org/10.1074/jbc.272.48.30061

73. Wysocki, R., Chéry, C. C., Wawrzycka, D., van Hulle, M., Cornelis, R., Thevelein, J. M., & Tamás, M. J. (2001). The glycerol channel Fps1p mediates the uptake of arsenite and antimonite in Saccharomyces cerevisiae. Molecular Microbiology, 40(6), 1391–1401. https://doi.org/10.1046/j.1365-2958.2001.02485.x

74. Wysocki, R., & Tamás, M. J. (2010). How Saccharomyces cerevisiae copes with toxic metals and metalloids. FEMS Microbiology Reviews, 34(6), 925–951. https://doi.org/10.1111/j.1574-6976.2010.00217.x

75. Wysocki, R., & Tamás, M. J. (2011). Saccharomyces cerevisiae as a Model Organism for Elucidating Arsenic Tolerance Mechanisms. In G. Banfalvi (Ed.), Cellular Effects of Heavy Metals (pp. 87–112). Springer Netherlands. https://doi.org/10.1007/978-94-007-0428-2_4

76. Yue, J.-X., Li, J., Aigrain, L., Hallin, J., Persson, K., Oliver, K., Bergström, A., Coupland, P., Warringer, J., Lagomarsino, M. C., Fischer, G., Durbin, R., & Liti, G. (2017). Contrasting evolutionary genome dynamics between domesticated and wild yeasts. Nature Genetics, 49(6), 913–924. https://doi.org/10.1038/ng.3847

77. Zabinsky, R. A., Mason, G. A., Queitsch, C., & Jarosz, D. F. (2019). It’s not magic – Hsp90 and its effects on genetic and epigenetic variation. Seminars in Cell and Developmental Biology, 88, 21–35. https://doi.org/10.1016/j.semcdb.2018.05.015

78. Zackrisson, M., Hallin, J., Ottosson, L.-G., Dahl, P., Fernandez-Parada, E., Ländström, E., Fernandez-Ricaud, L., Kaferle, P., Skyman, A., Stenberg, S., Omholt, S., Petrovič, U., Warringer, J., & Blomberg, A. (2016). Scan-o-matic: High-Resolution Microbial Phenomics at a Massive Scale. G3 Genes|Genomes|Genetics, 6(9), 3003–3014. https://doi.org/10.1534/g3.116.032342

