## Supplementary material for "Adaptation of the yeast gene knockout collection is near-perfectly predicted by fitness and diminishing return epistasis": Readme.txt

Supplementary data tables for “Adaptation of the yeast gene knockout collection is near-perfectly predicted by fitness and diminishing return epistasis”
Authors: Karl Persson, Simon Stenberg, Markus J. Tamás, Jonas Warringer

# Short Description:
Scan-o-matic growth phenotype datasets of the haploid BY4741 single gene deletion collection (MATa; his3Δ1; leu2Δ0; met15Δ0; ura3Δ0; GeneX::kanMX) adaptation to arsenite stress. From each growth curve, the colony populations doubling times (hours) have been extracted as a proxy for fitness (Zackrisson et al., 2016). 
Data S1: reports dose-response growth data for gene deletion strains exposed to different degrees of arsenite stress. Data S2: reports growth data, before LOESS fitting, for the complete collection of BY4741 gene deletion strains during adaptive evolution in 3 mM of arsenite. Data S3: reports growth data, after LOESS fitting, for the complete collection of BY4741 gene deletion strains after 25, 50 and 75 generations of adaptive evolution in 3 mM of arsenite. Data S4: reports growth data, before LOESS fitting, of BY4741 gene deletion strains during adaptive evolution across a range of selection pressures. Data S5: reports growth data, after LOESS fitting, of BY4741 gene deletion strains after 25, 50 and 75 generations across a range of selection pressures. Data S6: reports growth data, of BY4741 gene deletion strains on background media before and after adaptation to a range of selection pressures. Data S7: reports growth data, of single and double mutation strains in Synthetic complete medium, and arsenite, 3 mM.

# Column content
Tables are tab-separated values, where the doubling time phenotype is presented in hours for all datasets. Column content descriptions listed below:
S1: NaAsO2.mM [Concentration of arsenite], Gene.Systematic [Systematic gene name], Gene.Standard [Standard gene name], Doubling.time [Population doubling time]
S2: Gene.Standard [Standard gene name], Gene.Systematic [Systematic gene name], Cycle [Adaptation cycle], Generations [Generations passed on selective environment], Environment [Stress inducing environment], Doubling.time [Population doubling time], Fitted.Doubling.time [LOESS fitted Population doubling time], Strain.ID [Individual population ID: Gene.Standard-Plate-Row-Column]
S3: Strain.ID [Individual population ID: Gene.Standard-Plate-Row-Column], Doubling.time.first [Doubling time at first measurement], Doubling.time.25 [Doubling time at 25 generations], Doubling.time.50 [Doubling time at 50 generations], Doubling.time.75 [Doubling time at 75 generations], Doubling.time.last [Doubling time at final measurement], Gene.Systematic [Systematic gene name], Gene.Standard [Standard gene name]
S4: Gene.Standard [Standard gene name], Gene.Systematic [Systematic gene name], Cycle [Adaptation cycle], Generations [Generations passed on selective environment], Environment [Stress inducing environment], Doubling.time [Population doubling time], Fitted.Doubling.time [LOESS fitted Population doubling time], Strain.ID [Individual population ID: Environment-Gene.Standard-Plate-Row-Column]
S5: Strain.ID [Individual population ID: Environment-Gene.Standard-Plate-Row-Column], Doubling.time.first [Doubling time at first measurement], Doubling.time.25 [Doubling time at 25 generations], Doubling.time.50 [Doubling time at 50 generations], Doubling.time.75 [Doubling time at 75 generations], Doubling.time.last [Doubling time at final measurement], Gene.Systematic [Systematic gene name], Gene.Standard [Standard gene name], Environment [Stress inducing environment]
S6: Gene.Systematic [Systematic gene name], Environment [Stress inducing environment], Doubling.time.SC.before [Doubling time in background media before adaptation], Doubling.time.SC.after [Doubling time in background media after adaptation]
S7: Environment [Stress inducing environment], Gene.Standard [Standard gene name], Gene.Systematic [Systematic gene name], Secondary.Mutation [Secondary mutation on deletion mutant], Doubling.time.double.mutant [Doubling time of double mutant], Doubling.time.single.mutant [Doubling time with only the first mutation]

# Cultivation media
Solid media plates were cast with 50 ml of Synthetic Complete (SC) medium: 0.14% YeastNitrogenBase (YNB,CYN2210,ForMedium), 0.50% ammonium sulfate, 0.077% Complete Supplement Mixture (CSM, DCS0019, ForMedium), 2.0% (w/v) glucose and pH buffered to 5.8 with 1.0% (w/v) succinic acid, and 0.6% (w/v) NaOH and 20 g/L agar. Sterile filtered stress environments were supplemented to their indicated final concentrations. 

# References
Zackrisson, M., Hallin, J., Ottosson, L.-G., Dahl, P., Fernandez-Parada, E., Ländström, E., Fernandez-Ricaud, L., Kaferle, P., Skyman, A., Stenberg, S., Omholt, S., Petrovič, U., Warringer, J., & Blomberg, A. (2016). Scan-o-matic: High-Resolution Microbial Phenomics at a Massive Scale. G3 Genes|Genomes|Genetics, 6(9), 3003–3014. https://doi.org/10.1534/g3.116.032342
